# Genome-wide analysis identifies genetic effects on reproductive success and ongoing natural selection at the *FADS* locus

**DOI:** 10.1101/2020.05.19.104455

**Authors:** Iain Mathieson, Felix R. Day, Nicola Barban, Felix C. Tropf, David M. Brazel, eQTLGen Consortium, BIOS Consortium, Ahmad Vaez, Natalie van Zuydam, Bárbara D. Bitarello, Harold Snieder, Marcel den Hoed, Ken K. Ong, Melinda C. Mills, John R.B. Perry, on behalf of the Human Reproductive Behaviour Consortium

## Abstract

Identifying genetic determinants of reproductive success may highlight mechanisms underlying fertility and also identify alleles under present-day selection. Using data in 785,604 individuals of European ancestry, we identify 43 genomic loci associated with either number of children ever born (NEB) or childlessness. These loci span diverse aspects of reproductive biology across the life course, including puberty timing, age at first birth, sex hormone regulation and age at menopause. Missense alleles in *ARHGAP27* were associated with increased NEB but reduced reproductive lifespan, suggesting a trade-off between reproductive ageing and intensity. As NEB is one component of evolutionary fitness, our identified associations indicate loci under present-day natural selection. Accordingly, we find that NEB-increasing alleles have increased in frequency over the past two generations. Furthermore, integration with data from ancient selection scans identifies a unique example of an allele—*FADS1/2* gene locus—that has been under selection for thousands of years and remains under selection today. Collectively, our findings demonstrate that diverse biological mechanisms contribute to reproductive success, implicating both neuro-endocrine and behavioural influences.

## Introduction

Variation in human reproductive behaviour and success is epidemiologically associated with disease risk and has profound psychological, clinical, societal and economic implications. This is particularly true for infertility, where efforts to elucidate the underlying biological mechanisms have been limited by the lack of large, well-phenotyped studies with relevant infertility outcomes. This situation is mirrored across many reproductive traits and diseases, such as polycystic ovary syndrome^1,2^, where progress to identify genetic determinants and underlying mechanisms has lagged behind that of other complex diseases^3^.

One reason for this is that natural selection limits the frequency of fertility-reducing alleles. Number of children ever born (NEB) has one of the highest degrees of polygenicity of any complex trait, consistent with a genetic architecture strongly influenced by negative selection^4^. Studying the genetic basis of *fertility* may illuminate biological mechanisms underpinning infertility, with the advantage that relevant measures are more readily available. For example, recent studies have identified genetic determinants for NEB, age at first sexual intercourse and age at first birth^5–7^. These have provided several aetiological insights, such as highlighting a neuro-behavioural role for the estrogen receptor in men^5^ and identifying biological mechanisms linking reproductive ageing to late-onset diseases^5,6,8,9^.

Fertility-associated loci may act through a broad array of mechanisms. They may have direct effects on reproductive biology, or act through traits that contribute to partner selection or other aspects of behaviour and personality. For example, alleles associated with higher educational attainment are associated with lower fertility in some populations^10,11^, reflecting the link between higher education and older age at childbearing^12^. Finally, fertility-associated loci might represent alleles under selection for some trait entirely disconnected from reproductive biology. By definition, any variant that is under natural selection affects fitness. In particular, variants that affect fitness through NEB would be detected by a genome-wide scan for NEB, although this scan would not capture all components of fitness.

Our present study substantially builds upon two earlier studies^5,6^ to identify individual genetic determinants of NEB. We highlight a number of novel biological mechanisms and identify a locus that is under both historical and ongoing selection.

## Results

We identified genetic determinants of NEB by performing a genome-wide association study (GWAS) comprised of 785,604 European ancestry individuals (**Online Methods**) meta-analysed across 45 studies (**Table S1-S6**). SNP array data was imputed to at least 1000 Genomes Project reference panel density across all studies. The distribution of genome-wide test statistics for NEB showed substantial inflation (λ_GC_ = 1.36), however LD score regression^13^ indicated that this was attributable to polygenicity rather than population stratification (LD intercept 1.01; s.e. 0.008). In total, 5,283 variants reached genome-wide significance (P<5×10^-8^) for association with NEB, which we resolved to 28 statistically independent signals (**Table S7**). These include all six signals previously reported for NEB in overlapping samples of up to 343,072 individuals^5,6^.

The genetic architecture of NEB was only moderately correlated between men and women (r_g_=0.74; 95% CI 0.66-0.82). Therefore, we ran separate GWAS in men (N=306,980) and women (N=478,624), and identified six additional statistically independent signals (two in men, four in women). We found evidence of heterogeneity (P_het_<0.05) between sexes at 13/34 NEB loci (greater than expected by chance P_binomial_=4×10^-9^) and an overall trend for larger effect estimates in women than men (24/34, P_binomial_=0.02). Two notable examples were rs58117425 in testis expressed 41 (*TEX41*) gene which was significant only in men, and 6:152202621_GT_G in the estrogen receptor alpha (*ESR1*) where the effect on NEB in women was double that in men (**Table S7**).

In the absence of well-powered studies of infertility, we performed a GWAS on lifetime childlessness (CL) in UK Biobank (N=450,082) and assessed the relevance of NEB associated loci on susceptibility to CL. Effects on CL were modest, with the largest effect at the rs201815280-*CADM2* locus (sex combined OR=1.05, 95% CI [1.04-1.06], P=6.8×10^-18^]). Interestingly, the genetic correlation between NEB and CL was less than perfect (r_g_= −0.90 [−0.88 to −0.92). Accordingly, of the 16 independent loci identified for CL, eight were distinct from the NEB signals (**Table S7)**. Sex-stratified analyses revealed one additional female-specific CL signal (rs7580304, *PPP3R1*, **Table S7**). Several loci exhibited more significant effects on CL than on NEB (**Figure S1**).

In summary, we identified 43 independent signals; 28 from the sex-combined NEB meta-analysis, six sex-specific NEB signals, eight additional sex-combined CL signals, and one sex-specific CL locus. To validate these findings, we performed a lookup of these signals in 34,367 women from the FinnGen study (**Methods**). Since NEB was not recorded for men in this study we only considered the 41 signals identified in sex-combined or female-specific analyses. Despite the small replication sample, 35/41 effects were directionally concordant– (binomial sign test P=5×10^-6^; **Table S7**).

### Implicated genes and biological mechanisms

We identified putatively functional genes across all 43 NEB/CL associated loci using a number of approaches: gene expression integration, non-synonymous variant mapping and text mining (**Online Methods, Tables S8-S15**). Results are summarised in **Tables S16-17** and **Figure S2**. All prioritized genes were more highly expressed at the protein level in central nervous system cell types than in female reproductive cell types^14^, except for *ESR1* and *ENO4* (i.e. glandular cells in endometrium and fallopian tube in the case of *ENO4*, **Figure S2**). *ENO4* is required for sperm motility and function as well as for male fertility in mice. It is required for normal assembly of the sperm fibrous sheath, and provides most of the enolase activity in sperm^15^.

To further ascertain which loci might directly implicate reproductive mechanisms, we integrated GWAS results from other reproductive traits (**Figure 2**). Surprisingly, the NEB-increasing missense allele (rs9730, p.Ala117Thr) in *ARHGAP27*, which encodes a Rho GTPase, a small family of molecules involved in axon guidance, was associated with a shorter reproductive lifespan: later age at menarche (P=1×10^-11^) and earlier menopause (P=2×10^-5^); additional associations were seen with earlier age at first birth (P=5.5×10^-8^), lower circulating testosterone concentrations in women (both bioavailable [P=2.1×10^-4^] and total [P=2.1×10^-3^)], but higher testosterone concentrations in men (both bioavailable [P=3.5×10^-4^] and total [P=1.9×10^-11^]). These associations suggest a life history strategy involving a shorter but more productive reproductive lifespan. Another NEB signal, rs4730673 near *MDFIC*, is correlated with the strongest reported GWAS signal for same-sex sexual behaviour^16^ (rs10261857; r^2^ = 0.74). Here, the NEB-increasing allele had a more significant effect on decreasing the likelihood of CL, and also decreased the likelihood of same-sex sexual behaviour (**Table S7**).

**Figure 1.**
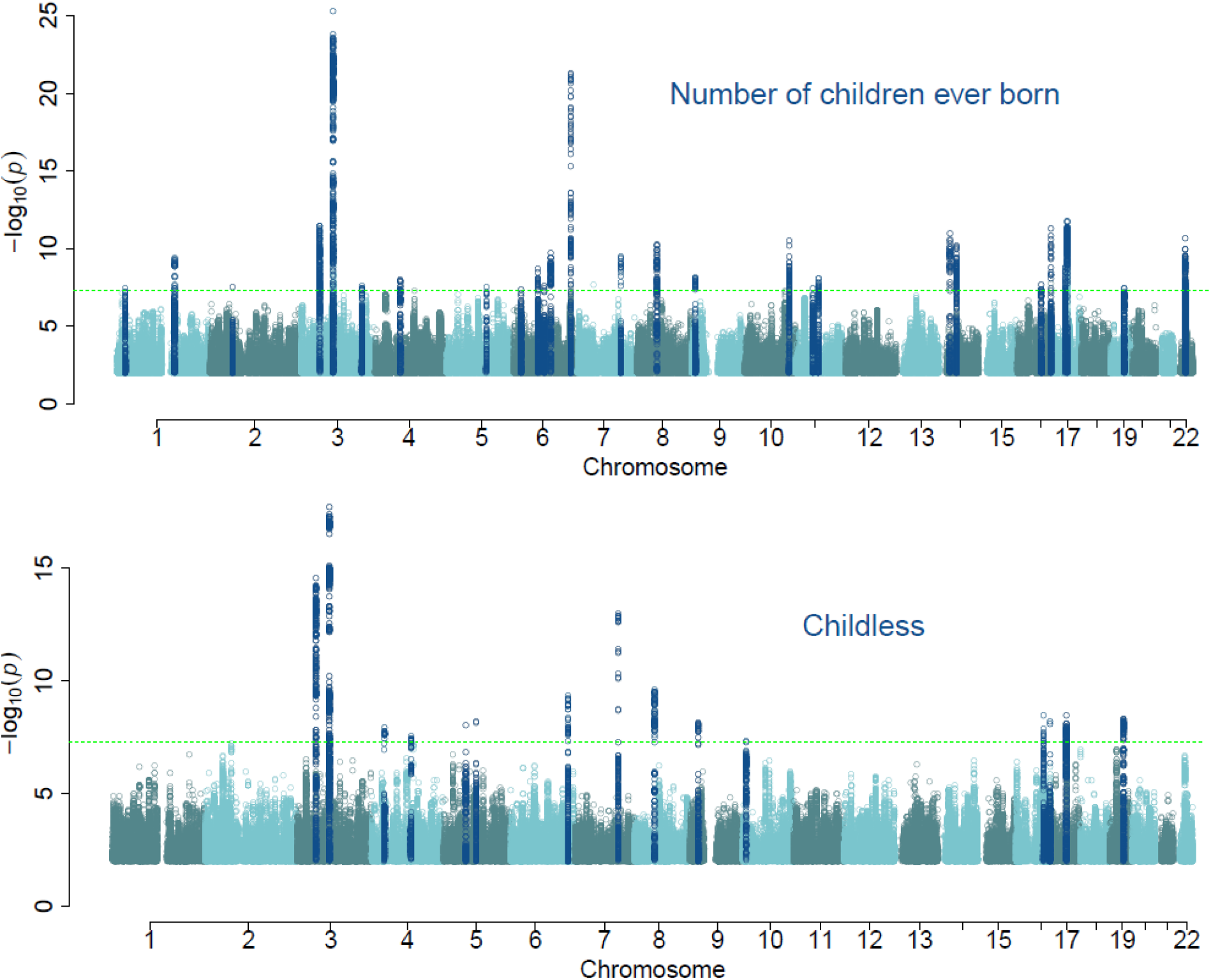
Manhattan plots for genome-wide association analyses of NEB and CL

**Figure 2.**
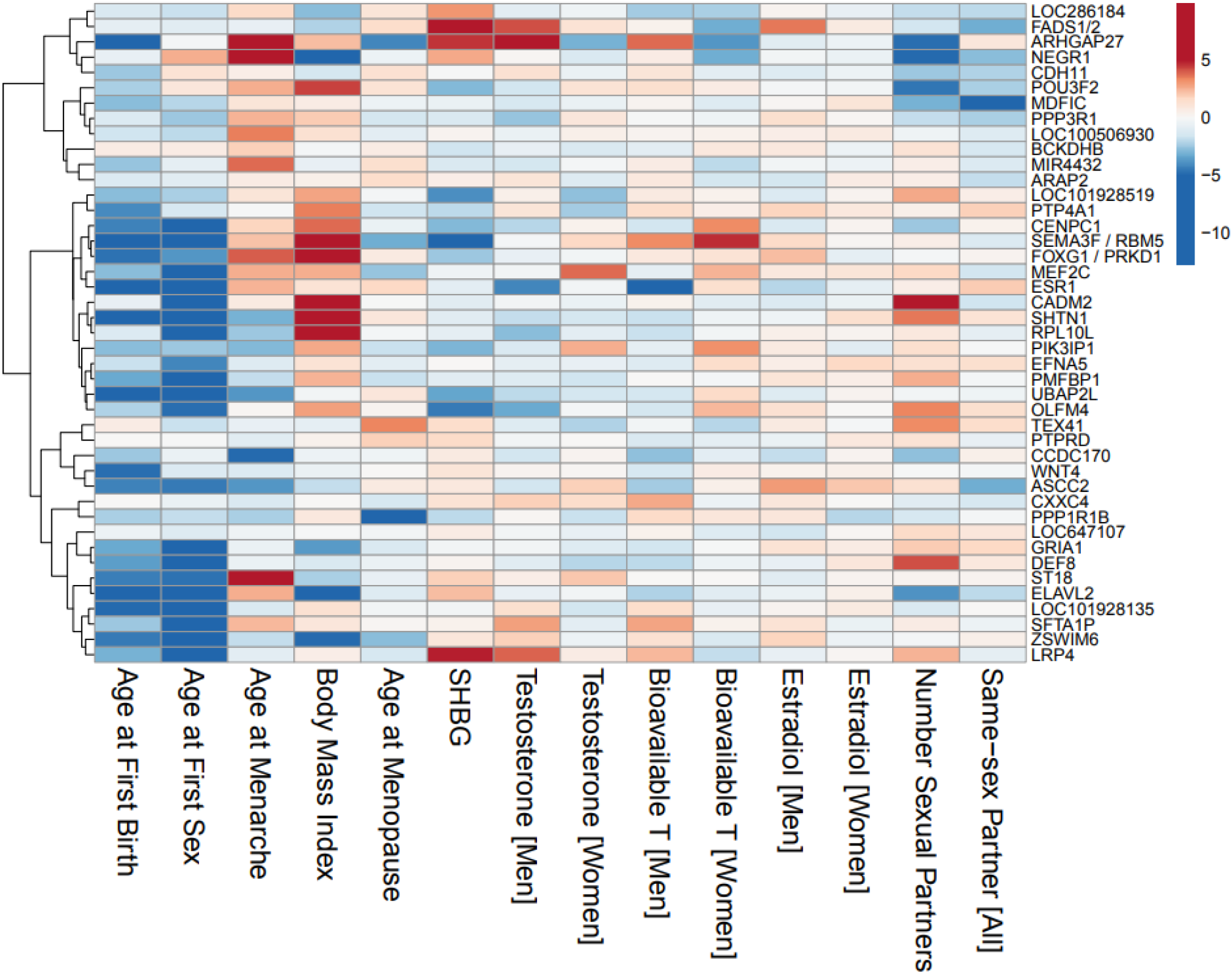
Heat map of the effects of the 43 independent signals identified for NEB or CL on other reproductive traits. All associations based on trait-specific Z-scores aligned to NEB-increasing allele, with 0 (white) denoting no association. SHBG = sex hormone binding globulin.

### NEB-increasing alleles have increased in frequency over the past two generations

We next tested whether NEB-increasing alleles have increased in frequency over time. To assess this, we computed a genome-wide polygenic score (PGS) for NEB using LDPred (**Online Methods**) and tested this as a linear function of birth year in studies independent of NEB PGS construction, controlling for the effects of study and sex. The standardised NEB PGS increased by 4.3×10^-3^ per-year (standard error = 6.4×10^-4^; P=1.3×10^-11^; **Figure S3**) in the HRS and UKHLS cohorts. This increase indicates that the identified loci are associated with increased NEB not only in our studied individuals, but also in their (unsampled) parents, since the PRS increasing over time suggests that parents with higher PRS have more children.

### Overlap between NEB and historical selection signals identifies the FADS1/2 locus

Effect estimates for the 34 genome-wide significant NEB loci ranged from 0.012-0.025 children per allele. The population mean NEB is ~1.8 in UK Biobank, so an effect size of 0.02 per allele implies that a group of 25 people homozygous for a NEB-increasing allele would have, on average, 46 children, compared to 45 children for a group of 25 people without that allele. Assuming no allelic effect on pre-reproductive mortality, these effects on NEB can be directly translated to selection coefficients of 0.67-1.4% per allele, which is within the range detectable by genome-wide historical selection scans^17–19^. Accordingly, we compared our NEB/CL GWAS results with the results of scans testing selection over different timescales from ~2,000 to ~30,000 years before present^17,18,20^ (**Online Methods**) and evaluated overlap using Bayesian co-localization analysis^21^ (**Table S18-S19**).

The strongest overlap was observed at chr11:61.5Mb, which exhibited a posterior probability of 96% that the lead variants for ancient selection and NEB represent the same underlying signal (**Figure 3A**). This locus contains the genes *FADS1* and *FADS2*, which have been targeted by selection multiple times in human history^22–25^. In particular, the derived haplotype at this locus has increased from a frequency of <10% 10,000 years ago to 60-75% in present-day European populations (**Figure 3B**). While some of this increase is due to admixture, there is strong evidence of positive selection over the past few thousand years, even accounting for changes in ancestry^17,22–26^, Each ‘C’ allele of the lead NEB SNP rs108499, which tags the positively selected *FADS1* haplotype, increased NEB by 0.0134, corresponding to a selection coefficient of 0.74% (0.0134 divided by mean NEB of 1.8).

**Figure 3.**
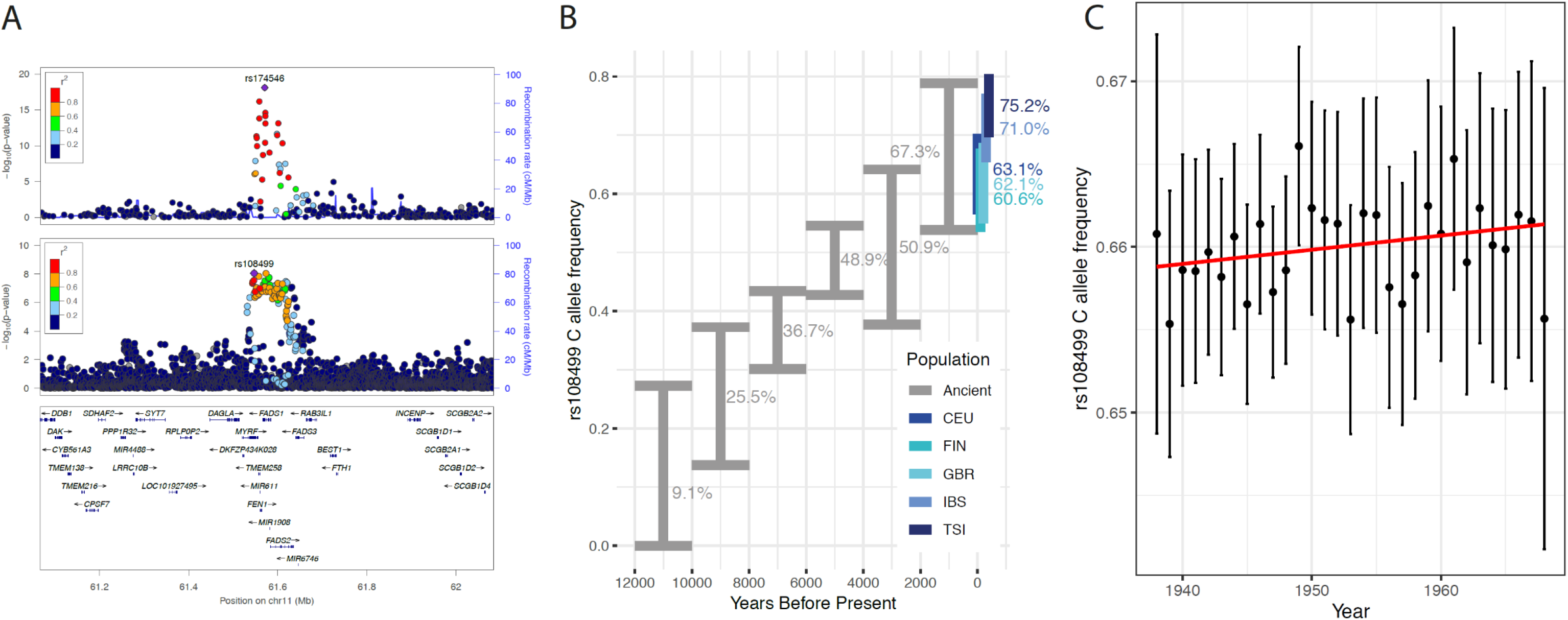
Evidence for historical and ongoing selection at the FADS locus. **A:** Colocalization of the ancient DNA selection signal^17^ (upper panel) and the NEB GWAS signal (lower panel). **B:** Estimated frequency (95% confidence intervals) for the derived *FADS* allele in Europe, based on direct evidence from ancient DNA. Present-day frequencies in 1000 Genomes European populations shown in blue. **C:** Frequency (95% confidence intervals) of the derived *FADS* allele in UK biobank as a function of birth year from 1938 to 1968.

Consistent with this, we estimate that within the “White British” subset of UK Biobank, the derived allele increased in frequency by 0.0088% per-year between the 1938 and 1969 birth cohorts (linear regression including 10 principal components and collection centre; P=0.27, **Figure 3C**) corresponding to a selection coefficient of 1.2% (approximate 95% CI −0.9-3.2%). Estimates of historical selection coefficients range from 0.4-0.6% (based on time-serial analysis of ancient DNA samples) to 3.4-6.9% (based on analyses of present-day haplotype structure)^24,26^.

*FADS1* and *FADS2* encode enzymes that catalyse the ω-5 and ω-6 lipid biosynthesis pathways that synthesize long chain polyunsaturated fatty acids (LC-PUFA) from short chain precursors. It has been hypothesised that the ancestral allele is advantageous in populations–for example in the Eurasian Upper Palaeolithic–that have a diet high in meat and marine fats (with relatively high LC-PUFA contents) and the derived allele is advantageous in populations that have a diet high in vegetable fats (with relatively low LC-PUFA contents and thus benefitting from higher FADS1 enzymatic activity)^22–25^. Under this model, FADS genotype selection in Europe is driven by dietary transitions, in particular, the Bronze Age transition to a diet based intensively on agricultural products^25,26^. However, the mechanism through which this gene-environment interaction affects fitness, and which phenotype is under selection, are unclear.

One reason for this uncertainty is that the *FADS* locus is highly pleiotropic. It is one of the strongest GWAS signals for circulating lipids^27^ and blood metabolites^28^, and is strongly associated with blood cell phenotypes, including erythrocyte and platelet sizes and counts (**Table S20**). To test the relevance of lipids to ongoing selection, we assessed the dose-response relationship of all previously reported^29^ HDL, LDL, total cholesterol or triglyceride associated variants on NEB using a Mendelian Randomization (MR) framework. We found no such association (P>0.05 in all MR models), which suggests that the NEB effect at the *FADS* locus is not shared across other lipid-associated loci. Assessment of the *FADS* locus across a range of reproductive traits highlighted associations between the NEB-increasing allele and higher circulating sex-hormone binding globulin (SHBG, P=2.3×10^-20^), higher total testosterone (P=1.9×10^-5^) and higher estradiol concentrations (P=4×10^-4^) in men, and lower bioavailable testosterone concentrations in women (P=1.5×10^-3^). These data, in addition to the observation that our associated variants influence *FADS1* expression in the brain (**Table S15**), raise the possibility that the effect of the *FADS* locus on NEB could be attributable to its effects on reproductive pathways, rather than its effects on lipids and consequent disease morbidity/mortality. We note however that many of the strongest known genetic determinants for sex hormone levels^30^ are not significantly associated with NEB in our study. Ultimately further experimental work is required to fully elucidate the mechanisms linking NEB-associated variants at this locus to reproductive success. We note that our analyses include only European ancestry individuals and is heavily weighted by the UK Biobank, which may not be representative of the UK population,^31^ and it remains to be seen which of these effects are consistent across cohorts and populations.

### NEB-associated variants in CADM2 exhibit signatures of balancing selection

The most significant NEB-associated variants in the genome, in *CADM2*, show no evidence of historical positive selection. However, *CADM2* is reported to exhibit one of the strongest genomic signals of long-term balancing selection^32^. Variants in *CADM2* are associated with a range of behavioural and reproductive traits, plausibly explained by a primary effect on risk taking propensity^5,33,34^ Variants that increase risk taking also increase NEB, with risk taking and behavioural disinhibition also linked to earlier reproductive onset ^7^. Since these variants are still segregating and appear to be under balancing rather than positive selection, this suggests ongoing or time- or environmentally-varying negative consequences of NEB-increasing *CADM2* alleles on fitness, possibly connected to adverse risk-taking behaviours. Other NEB-associated loci with nominal evidence of balancing selection contain the genes *PTPRD* and *LINC00871* (**Table S21**).

### Lack of contemporary selection at historical selection signals

We next tested whether there was any evidence of current selection (as measured by association with NEB/CL) at regions identified by the three genome-wide historical selection scans (**Table S22**). Other than the *FADS* locus, none of the other 53 regions tested exhibited an association with NEB, suggesting that few of the strong historical selective sweeps in humans are ongoing. For example, the sweep associated with lactase persistence - one of the strongest signals of selection in any human population – is not ongoing in the European ancestry populations in this study. This finding is consistent with evidence from ancient DNA which suggests that the persistence allele reached its present-day Northern European frequency of around 60% by the Middle Ages^35^. Other strong sweeps, such as those associated with skin pigmentation-decreasing loci, are likely not detected in the NEB GWAS because the selected variants are now fixed or almost fixed among European ancestry populations.

Different methods for detecting historical selection in humans are sensitive to selection across very different timescales – ranging from thousands to millions of years. Our NEB GWAS can be interpreted as a genome-wide selection scan over the shortest and most recent timescale – i.e., living generations. The limited overlap between this and the historical selection scans is consistent with the limited overlap between different historical scans, and likely reflects a highly dynamic landscape of selection. Selected loci either fix, or stop being selected and remain at intermediate frequency. The *FADS* locus is unique in the sense that the selective sweep – starting at least several thousand years ago – is still ongoing. At other loci, for example *CADM2*, signs of balancing selection suggest that the direction of effect might vary with environments or allele frequency.

In summary, our study identifies 37 signals that have not been previously reported for NEB and represent potential targets of ongoing natural selection. Further work should aim to parse these effects into mechanisms that directly influence reproductive biology, in contrast to those which affect behaviour or reduce fitness through premature morbidity or mortality.

## Acknowledgements

This research was conducted using the UK Biobank Resource under application 32696 and 9797. This work was supported by the Medical Research Council [Unit Programme number MC_UU_12015/2], ERC grants 615603 and 835079, NIH grant R35GM133708 and The Leverhulme Trust, Leverhulme Centre for Demographic Science. Full study-specific and individual acknowledgements can be found in the supplementary information. The content is solely the responsibility of the authors and does not necessarily represent the official views of the National Institutes of Health or other funders.

## Online Methods

### Phenotype definitions

*Number of children ever born (NEB)* is treated as a continuous measure that was either asked directly or could be created from several survey questions (e.g., pregnancy or birth histories). A standard question in most surveys asks: *How many children have you given birth to?* Or another variant is*: How many children do you have?* In most cases it was possible to distinguish between biological, adopted or step-children and when this was possible, we refer to live born biological children. Individuals were eligible for inclusion in the analysis if they were assessed for NEB and were at least age 45 for women and age 55 for men, or in other words, had reached the end of their reproductive window. The measure included both those who had given birth to a child (parous) and those who had not (nulliparous).

*Childlessness (CL)* is treated as a binary measure, generally calculated from NEB recoded as 1 referring to childless and 0 if they had children with the same inclusion rules of biological live born children and age restrictions applied. Detailed measures for both phenotypes per cohort are described in **Table S2**).

### Participating cohorts and analysis plan

The discovery of genetic variants for reproductive success in human populations is based on genome-wide association studies from cohort-level data that were quality-controlled and meta-analyzed by two separate independent centres at the University of Oxford and University of Cambridge. We follow the QC protocol of the GIANT consortium’s study of human height^36^ and employed the software packages QCGWAS^37^ and EasyQC^38^, which allowed us to harmonize the files and identify possible sources of errors in association results. This procedure entailed that diagnostic graphs and statistics were generated for each set of GWAS results (i.e., for each file). In the case where apparent errors could not be amended by stringent QC and correspondence with the local analyst of the respective cohort, cohorts were excluded from the meta-analysis. (See section below for details on cohort inclusion and errors).

A total of 45 cohorts participated in our study (**Table S1**). **Table S2** provides an overview of cohort-specific details, including an adjusted pooled analysis of women and men in the case of family data (see below). Cohorts who agreed to participate followed an Analysis Plan posted on the Open Science Framework preregistration site https://osf.io/b4r4b/ on February 08, 2017.

For autosomal chromosomes and NEB the total number of individuals in the pooled metaanalysis was 785,604, with a somewhat larger sample for women (39 cohorts) than men (29 cohorts). **Table S3** provides detailed information about the cohorts with X Chromosome data. For the NEB meta-analysis this included 671,349 individuals (12 cohorts) for the pooled cohorts, 384,976 for women (12 cohorts) and 286,373 for men (10 cohorts). For CL, only data from the UK Biobank was used, with 450,082 individuals in the pooled analysis, 245,047 for women and 205,035 for men – both for autosomal chromosomes and X chromosome.

### Sample exclusion criteria

Individuals were eligible for inclusion if they met the following conditions:

a. They were assessed for NEB at least at age 45 for women, age 55 for men;
b. Those who have both given birth to a child (parous) and those who have not (nulliparous);
c. All relevant covariates (e. g. year of birth) are available for the individual;
d. They were successfully genotyped genome-wide (recommended individual genotyping rate > 95%);
e. They passed the cohort-specific standard quality controls, e.g. excluding individuals who are genetic outliers in the cohort.
f. They were of European ancestry.

### Genotyping and Imputation

**Table S4** provides an overview of the cohort-specific details on the genotyping platform, pre-imputation quality control filters applied to the genotype data, imputation software used, the reference used for imputation and the presence of X chromosome data. We asked cohorts to include all autosomal SNPs imputed from the 1000G panel (at a minimum) to allow analyses across different genotyping platforms. Cohorts with denser reference panels we asked to communicate this to our team. Cohorts were asked to provide unfiltered results since filters on imputed markers and so forth would be applied at the meta-analysis stage.

### Association testing models

Analysts ran linear regression models for NEB and logistic regression models for CL in the UK Biobank study. Analysts were asked to include birth year of the respondent (represented by birth year minus 1900), its square and cubic to control for non-linear birth cohort effects. For those with family based data, we suggested controlling for family structure or excluding relatives. We furthermore asked studies with family data to run a pooled GWAS on both sexes. Combined analyses that included both men and women also needed to include interactions of birth year and its polynomials with sex. We asked cohorts to include top principal components to control for population stratification^39^ and cohort specific covariates if appropriate. Some cohorts only used birth year and not the polynomials because of multi-collinearity issues/convergence of the GWA analysis.

### Analysis of X chromosome

Analysis of X chromosome variants was performed using one of two complementary approaches, XWAS or SNPtest, the results of which could be combined by meta-analysis. In XWAS software (http://keinanlab.cb.bscb.cornell.edu/content/xwas) we used the *--var-het-weight* command. In SNPtest, we used the -method *newml*; while this assumes complete X-inactivation (i.e. a male with one allele is considered the same as a homozygous female) the effect estimates and standard errors approximate ½ of those produced by XWAS.

### Quality Control: filters & diagnostic checks

We followed the quality control (QC) protocol reported by the GIANT consortium’s GWAS of height^36^. We used an adapted version of the software package QCGWAS^37^, which allows the inclusion of structural variants, in order to standardize files across cohorts and we used EasyQC^38^ to filter variants by QC criteria and to produce diagnostic graphs and statistics as described below. Where errors could not be amended by combining stringent QC with file-inspections, queries to cohorts and corrections, cohorts were excluded from the meta-analysis. See also Supplementary **Tables S5-S6** for QC results on autosomal and X chromosomes for NEB and CL. Specific individual filters were:

a. **Missing data.** We filtered variants where information on both reference and other allele were missing, where the estimated effect, *p*-value, standard error, expected allele frequency or number of observations were missing.
b. **Implausible values.** We filtered variants where *p*-values > 1 or < 0, standard errors = 0 or = infinite, expected allele frequency > 1 or < 0, N < 0, call rate > 1 or < 0, an SE of the effect estimate which is approximately 40% greater than the expected SE based on MAF and standard deviation and for those with an *R^2^* >10% (see Winkler et al^38^ for an the approximation for quantitative and Rietveld et al^40^ for quantitative and binary traits).
c. **Quality thresholds.** We filtered variants where expected allele frequency = 1 or = 0 (monomorphic variants), N < 100 to guard against spurious associations due to overfitting of the model, minor allele count <6 to guard against spurious associations with low frequency-SNPs and genotyped SNPs which were not in Hardy-Weinberg Equilibrium (HWE), with significant thresholds of 10^-3^ in case N < 1,000, 10^-4^ in case 1,000 ≤ N < 2,000, 10^-5^ in case 2,000≤N<10,000 and no filter in case N>10,000, imputed markers with imputation quality < 40% and SNPs with a call rate < 95%, if discrepancies between reported and expected *p*-value based on effect estimates and standard errors are detected (see also next section on diagnostic graphs).
d. **Data harmonization.** We matched the cohort based summary statistics with a 1000 Genome reference panel phase 1 version 3 reference panel provided by Winkler et al^38^. EasyQC drops mismatched variants which cannot be solved straight away such as duplicates, allele mismatches or missing or invalid alleles. Based on graphical inspections (see below), we applied cohort specific filters to drop variants with obvious deviations between expected allele frequency based on the reference panel and observed allele frequency.

### Quality Control: Diagnostic graphs

We produced three key diagnostic graphs for visual inspection by the two independent QC centres in Oxford and Cambridge. If problems were detected which could not be resolved by more stringent QC we had to remove the cohort from the analysis. The key diagnostic graphs depicted:

a. An **allele frequency (AF) plot** to identify errors in allele frequencies and strand orientations using the 1000 Genome phase 1 version 3 reference panel provided by Winkler et al^38^.
b. A **PZ plot** to assess the consistency of the reported p-values versus the Z score calculated based on effect sizes and standard errors.
c. A **PRS plot** of predicted versus reported standard error as developed by Winkler et al^38^ and implemented by Okbay et al^41^.

### Quality Control: Filtering results

a. Autosomal chromosomes Overall, the quality of studies was good (for full results of the QC-filters described above see **Tables S5** and **S6** for autosomal SNPs). One cohort needed to be excluded (INGI-Carlantino) due to the filter on sample size. For autosomal chromosomes and NEB, the remaining 45 cohorts provided 81 files, 39 for women only, 28 for men only and 13 pooled (for family data). Two studies did not provide imputation quality (KORA F3, N =1.066; and KORA F4, N =1,111) and contributed only 584,866 and 496,556 SNPs respectively. For the two HPFS cohorts, results from our last discovery GWAS^6^ based on HapMap 2 reference panels were recycled with number of SNPs between 2,394,353 and 2,412,487. For all other cohorts, the number of variants in the analysis range between 6,691,978 for men in LBC 1921 and 20,783,286 for women in EPIC with an average of 10,574,721. For CL, between 25,555,939 and 25,554,098 variants from the UK Biobank entered the GWAS and between 13,539,540 and 13,661,642 survived the QC.
b. X chromosome For NEB, 12 cohorts provided information on the X chromosome. Overall, we received 27 files, 12 for women, 10 for men and 5 for the pooled analysis in case there were relatives in the data. On average 325,872 variants survived QC with a minimum of 191,880 in men from LBC 1921 to 991,081 for the pooled UK Biobank sample. For CL, the UK Biobank provided results for between 980,779 and 991,081 variants on the X chromosome after QC.

### GWAS meta-analysis, signal selection and replication

Cohort association results (after applying the QC filters) were combined using sample-size weighted meta-analysis, implemented in METAL^42^. Sample-size weighting is based on Z-scores and can account for different phenotypic measurements among cohorts^43^. The two QC centres agreed in using sample-size weighting to allow cohorts to introduce study-specific covariates in their cohort-level analysis. Only SNPs that were observed in at least 50% of the participants for a given phenotype-sex combination were passed to the meta-analysis. SNPs were considered genome-wide significant at *p*-values smaller than 5×10-8 (α of 5%, Bonferroni-corrected for a million tests). The meta-analyses were carried out by two independent analysts. Comparisons were made to ensure concordance of the identified signals between the two independent analysts. Distance-based clumping (using a 1Mb window) was used to identify the most significant SNPs in associated regions (termed “lead SNPs”). This was then supplemented by approximate conditional analysis implemented in GCTA^44,45^, where we required additional signals to be genome-wide significant in both pre and post conditional models.

We meta-analysed GWAS results for NEB and CL both in sex-combined and sex-specific models. To understand the magnitude of the estimated effects, we used an approximation method to compute unstandardized regression coefficients based on the Z-scores of METAL output obtained by sample-size-weighted meta-analysis, allele frequency and phenotype standard deviation.

### Replication

Replication was performed using the FinnGen study - a public-private partnership project combining genotype data from Finnish biobanks and digital health record data from Finnish health registries (https://www.finngen.fi/en). Six regional and three country-wide Finnish biobanks participate in FinnGen. Additionally, data from previously established populations and disease-based cohorts are utilized. Release 4 of Finngen includes 176,899 participants.

In this analysis we included women that participated to Finngen release 4 and were at least 45 years of age at 31^st^ December 2017. This was the last date we had information from national registries. We also excluded women younger than 16 in 1969 (the start of the registries). Using these inclusion criteria, we included women born between 1953 and 1973 and children born between 1969 and 2017. We also excluded women that emigrated from Finland in the study period.

To determine if a woman delivered a child we used the following codes obtains from the national inpatient registry (HILMO):

- ICD-10 codes: O80-O84
- ICD-9 code: 6440B, 6450B, 650[0-9]B-659[0-9]B
- ICD-8 codes: 650-662

When multiple codes were used within a 10-month period we counted as a single delivery.

There were 37,741 women, the average (SD) number of children was 1.72 (1.32) and 20.4% of the women were childless.

Samples were genotyped with Illumina (Illumina Inc., San Diego, CA, USA) and Affymetrix arrays (Thermo Fisher Scientific, Santa Clara, CA, USA). Genotype calls were made with GenCall and zCall algorithms for Illumina and AxiomGT1 algorithm for Affymetrix data. Chip genotyping data produced with previous chip platforms and reference genome builds were lifted over to build version 38 (GRCh38/hg38) following the protocol described here: dx.doi.org/10.17504/protocols.io.nqtddwn. In sample-wise quality control, individuals with ambiguous sex, high genotype missingness (>5%), excess heterozygosity (+-4SD) and non-Finnish ancestry were removed. In variant-wise quality control variants with high missingness (>2%), low HWE P-value (<1e-6) and minor allele count, MAC<3 were removed. Chip genotyped samples were pre-phased with Eagle 2.3.5 (https://data.broadinstitute.org/alkesgroup/Eagle/) with the default parameters, except the number of conditioning haplotypes was set to 20,000.

High-coverage (25-30x) WGS data (N= 3,775) were generated at the Broad Institute and at the McDonnell Genome Institute at Washington University; and jointly processed at the Broad Institute. Variant call set was produced with GATK HaplotypeCaller algorithm by following GATK best-practices for variant calling. Genotype-, sample- and variant-wise QC was applied in an iterative manner by using the Hail framework (https://github.com/hail-is/hail) v0.1 and the resulting high-quality WGS data for 3,775 individuals were phased with Eagle 2.3.5 as described above. Genotype imputation was carried out by using the population-specific SISu v3 imputation reference panel with Beagle 4.1 (version 08Jun17.d8b, https://faculty.washington.edu/browning/beagle/b4_1.html) as described in the following protocol: dx.doi.org/10.17504/protocols.io.nmndc5e. Post-imputation quality-control involved non-reference concordance analyses, checking expected conformity of the imputation INFO-values distribution, MAF differences between the target dataset and the imputation reference panel and checking chromosomal continuity of the imputed genotype calls.

For principal components analysis, FinnGen data was combined with 1000 genomes data. Related individuals (<3rd degree) were removed using King software^46^. We considered common (MAF >= 0.05) high quality variants: not in chromosome X, imputation INFO>0.95, genotype imputed posterior probability>0.95 and missingess<0.01. LD-pruned (r^2^<0.1) common variants were used for computing PCA with Plink 1.92.

SAIGE mixed model logistic regression (https://github.com/weizhouUMICH/SAIGE/releases/tag/0.35.8.8) was used for association analysis. Age and 10 PCs and genotyping batch were used as covariates. Each genotyping batch was included as a covariate to avoid convergence issues.

### Prioritizing putatively functional genes in GWAS highlighted regions

We used a range of procedures to prioritize the most likely causal gene(s) in NEB and CLN-associated loci before combining results across the two traits.

Firstly, we used DEPICT v1.1^47^ to identify enrichment for pathways, cell types and tissues, as well as to prioritize candidate genes. DEPICT is an integrative tool that employs predicted gene functions and that is agnostic to the outcomes analysed in the GWAS. For both NEB and CLN, all SNPs with *P*<1×10^-5^ in the pooled analyses were used as input.

Secondly, we used Phenolyzer (v1.1), to prioritize candidate genes by integrating prior knowledge and phenotype information^48^. Here we used the regions defined by DEPICT (see above), reflecting loci reaching *P*<1×10^-5^ in first instance. Phenolyzer takes free text input and interprets these as disease names by using a word cloud to identify synonyms. It then queries precompiled databases for the disease names to find and score relevant seed genes. The seed genes are subsequently expanded to include related (predicted) genes based on several types of relationships, e.g. protein-protein interactions, transcriptional regulation and biological pathways. Phenolyzer uses machine learning techniques on seed genes and predicted gene rankings to produce an integrated score for each gene. We used search terms capturing three broad areas, i.e. (in)fertility, congenital neurological disorders and psychological traits, based on results from pathway, tissue and cell type enrichment analyses. Phenolyzer identified 16 and 8 candidate genes for NEB and CLN, respectively for gene scores >0.1 within the selected DEPICT regions with genome-wide significant lead SNP. Where more than one gene was prioritized per locus the gene with the highest total score was reported.

Thirdly, we used *in silico* sequencing to identify non-synonymous variants with an R^2^ for LD>0.7 with the lead SNPs of NEB and CL-associated loci.

Finally, we used Summary-data-based Mendelian Randomization (SMR) and heterogeneity in dependent instruments (HEIDI)^49^ using data from the eQTL consortium in whole blood^50^, and a meta-analysis of eQTL data in brain^51^.

We integrated findings across all approaches described above and retained genes in loci that reached genome-wide significance and that were located within 1Mb from a GWAS lead SNP.

### Testing change in polygenic scores over time

To evaluate how NEB-associated alleles may change in frequency over time, we first repeated our NEB GWAS meta-analysis twice excluding either the Health and Retirement Study (HRS) and the UK Household Longitudinal Study - Understanding Society (UKHLS). Polygenic risk scores were then generated from the meta-analysis excluding either HRS or UKHLS and used to predict outcomes in the respective study. Polygenic risk scores were computed using LDPred^52^ from the meta-analyses, using the LD reference panel from the respective target studies (HRS or UKHLS). We calculated LDpred weights under the infinitesimal model, using summary statistics from the sex-combined meta-analysis results excluding the specific cohort from the calculation. We then performed a linear regression of the polygenic scores for males and females separately against birth year, controlling for the first 10 genomic principal components. We excluded birth years with fewer than 90 individuals.

### Identifying overlap between NEB hits and previously-identified selection signals

We deployed several methods to assess overlap of our NEB signals with three genome-wide selection scans. First, the Composite of Multiple Signals test^20^ combines information from different statistics to detect selection on the order of the past 50,000 years. Second, an ancient DNA based scan^17^ that uses direct inference of allele frequency from ancient populations to infer selection over the past 10,000 years. Finally, the Singleton Density Score^18^, which uses patterns of singleton variants to infer very recent selection – on the order of a few thousand years.

We obtained genome-wide selection scan results from three sources^17,18,20^. For the Composite of Multiple Signals test^20^, we used the rankings of *CMS_GW_* statistics to obtain an empirical P-value for each SNP. For the Singleton Density Score ^18^, we converted normalized SDS scores to two-tailed P-values of the standard normal distribution. Finally, for the ancient DNA based selection scan^17^, we used the genomic control corrected P-values from the original scan. For each NEB hit, for each scan, we identified the SNP within 10kb, with a P-value < 10^-6^ for NEB that had the most significant selection scan signal (SNP1 and PVAL1 in **Table S18-19**). We also identified the SNP within 10kb that had the most significant selection scan signal, regardless of its NEB P-value (SNP2 and PVAL2 in **Table S18-19**).

Finally, we performed a Bayesian Colocalization analysis using the “*coloc*” package^21^ using all SNPs within 10kb of the lead NEB SNP. This computes posterior probabilities for the hypotheses: H0 No causal SNPs, H1 Causal SNP for selection but not NEB, H2 Causal SNP for NEB but not selection, H3 One independent causal SNP for each trait, H4 One shared causal SNP for both traits. We report the hypothesis with the maximum posterior probability (COLOC in **Table S18-19**).

We also tested for overlap with a scan for balancing selection using the NCD2 statistic^53^ for the GBR population of 1000 genomes. We used the target frequency of 0.5 for these tests. For all SNPs, we report the value of the window that overlaps that SNP or, if more than one window overlaps a SNP, we report the lowest P-value of any window within 10kb. Finally, we report the lowest P-value for genes within 10kb of each SNP.

### Estimating *FADS* allele frequencies from ancient DNA

*We* downloaded combined data from https://reich.hms.harvard.edu/downloadable-genotypes-present-day-and-ancient-dna-data-compiled-published-papers and restricted to 652 samples west of 40°E, north of 35°S, more recent than 12,000 years before present and with coverage at rs108499. We binned them into 2000-year bins, and computed estimated allele frequencies and bootstrap confidence intervals. We also include the European sub-populations from phase 3 of the 1000 Genomes Project^54^.

## References

1. Day, F. R. et al. Causal mechanisms and balancing selection inferred from genetic associations with polycystic ovary syndrome. Nat. Commun. 6, 8464 (2015).

2. Day, F. et al. Large-scale genome-wide meta-analysis of polycystic ovary syndrome suggests shared genetic architecture for different diagnosis criteria. PLoS Genet. 14, e1007813 (2018).

3. Censin, J. C., Bovijn, J., Holmes, M. V & Lindgren, C. M. Commentary: Mendelian randomization and women’s health. Int. J. Epidemiol. 48, 830–833 (2019).

4. O’Connor, L. J. et al. Extreme Polygenicity of Complex Traits Is Explained by Negative Selection. Am. J. Hum. Genet. 105, 456–476 (2019).

5. Day, F. R. et al. Physical and neurobehavioral determinants of reproductive onset and success. Nat. Genet. 48, 617–623 (2016).

6. Barban, N. et al. Genome-wide analysis identifies 12 loci influencing human reproductive behavior. Nat. Genet. 48, 1462–1472 (2016).

7. Mills, M. C. et al. Large-scale genome-wide association study identifies 370 loci for onset of sexual and reproductive behaviour implicated with psychiatric disorders, infertility and longevity. bioRxiv (2020) doi:10.1101/2020.05.06.081273.

8. Day, F. R. et al. Genomic analyses identify hundreds of variants associated with age at menarche and support a role for puberty timing in cancer risk. Nat. Genet. 10, 1–19 (2017).

9. Day, F. R. et al. Large-scale genomic analyses link reproductive aging to hypothalamic signaling, breast cancer susceptibility and BRCA1-mediated DNA repair. Nat. Genet. 47, 1294–303 (2015).

10. Kong, A. et al. Selection against variants in the genome associated with educational attainment. Proc. Natl. Acad. Sci. U. S. A. 114, E727–E732 (2017).

11. Beauchamp, J. P. Genetic evidence for natural selection in humans in the contemporary United States. Proc. Natl. Acad. Sci. U. S. A. 113, 7774–9 (2016).

12. Mills, M., Rindfuss, R. R., McDonald, P. & te Velde, E. Why do people postpone parenthood? Reasons and social policy incentives. Hum. Reprod. Update 17, 848–860 (2011).

13. Bulik-Sullivan, B. K. et al. LD Score regression distinguishes confounding from polygenicity in genome-wide association studies. Nat. Genet. 47, 291–5 (2015).

14. Uhlén, M. et al. Proteomics. Tissue-based map of the human proteome. Science 347, 1260419 (2015).

15. Nakamura, N. et al. Disruption of a spermatogenic cell-specific mouse enolase 4 (eno4) gene causes sperm structural defects and male infertility. Biol. Reprod. 88, 90 (2013).

16. Ganna, A. et al. Large-scale GWAS reveals insights into the genetic architecture of same-sex sexual behavior. Science 365, (2019).

17. Mathieson, I. et al. Genome-wide patterns of selection in 230 ancient Eurasians. Nature 528, 499–503 (2015).

18. Field, Y. et al. Detection of human adaptation during the past 2000 years. Science 354, 760–764 (2016).

19. Grossman, S. R. et al. A composite of multiple signals distinguishes causal variants in regions of positive selection. Science 327, 883–6 (2010).

20. Grossman, S. R. et al. Identifying recent adaptations in large-scale genomic data. Cell 152, 703–13 (2013).

21. Giambartolomei, C. et al. Bayesian test for colocalisation between pairs of genetic association studies using summary statistics. PLoS Genet. 10, e1004383 (2014).

22. Fumagalli, M. et al. Greenlandic Inuit show genetic signatures of diet and climate adaptation. Science 349, 1343–7 (2015).

23. Ameur, A. et al. Genetic adaptation of fatty-acid metabolism: a human-specific haplotype increasing the biosynthesis of long-chain omega-3 and omega-6 fatty acids. Am. J. Hum. Genet. 90, 809–20 (2012).

24. Ye, K., Gao, F., Wang, D., Bar-Yosef, O. & Keinan, A. Dietary adaptation of FADS genes in Europe varied across time and geography. Nat. Ecol. Evol. 1, 167 (2017).

25. Buckley, M. T. et al. Selection in Europeans on Fatty Acid Desaturases Associated with Dietary Changes. Mol. Biol. Evol. 34, 1307–1318 (2017).

26. Mathieson, S. & Mathieson, I. FADS1 and the Timing of Human Adaptation to Agriculture. Mol. Biol. Evol. 35, 2957–2970 (2018).

27. Teslovich, T. M. et al. Biological, clinical and population relevance of 95 loci for blood lipids. Nature 466, 707–13 (2010).

28. Draisma, H. H. M. et al. Genome-wide association study identifies novel genetic variants contributing to variation in blood metabolite levels. Nat. Commun. 6, (2015).

29. Klarin, D. et al. Genetics of blood lipids among ~300,000 multi-ethnic participants of the Million Veteran Program. Nat. Genet. 50, 1514–1523 (2018).

30. Ruth, K. S. et al. Using human genetics to understand the disease impacts of testosterone in men and women. Nat. Med. (2020).

31. Fry, A. et al. Comparison of Sociodemographic and Health-Related Characteristics of UK Biobank Participants With Those of the General Population. Am. J. Epidemiol. 186, 1026–1034 (2017).

32. Siewert, K. M. & Voight, B. F. Detecting Long-Term Balancing Selection Using Allele Frequency Correlation. Mol. Biol. Evol. 34, 2996–3005 (2017).

33. Boutwell, B. et al. Replication and characterization of CADM2 and MSRA genes on human behavior. Heliyon 3, e00349 (2017).

34. Day, F. R., Ong, K. K. & Perry, J. R. B. Elucidating the genetic basis of social interaction and isolation. Nat. Commun. 9, 2457 (2018).

35. Krüttli, A. et al. Ancient DNA analysis reveals high frequency of European lactase persistence allele (T-13910) in medieval central europe. PLoS One 9, e86251 (2014).

36. Wood, A. R. et al. Defining the role of common variation in the genomic and biological architecture of adult human height. Nat. Genet. 46, 1173–86 (2014).

37. van der Most, P. J. et al. QCGWAS: A flexible R package for automated quality control of genome-wide association results. Bioinformatics 30, 1185–1186 (2014).

38. Winkler, T. W. et al. Quality control and conduct of genome-wide association metaanalyses. Nat. Protoc. 9, 1192–212 (2014).

39. Price, A. L. et al. Principal components analysis corrects for stratification in genomewide association studies. Nat. Genet. 38, 904–9 (2006).

40. Rietveld, C. A. et al. GWAS of 126,559 individuals identifies genetic variants associated with educational attainment. Science 340, 1467–71 (2013).

41. Okbay, A. et al. Genome-wide association study identifies 74 loci associated with educational attainment. Nature 533, 539–42 (2016).

42. Willer, C. J., Li, Y. & Abecasis, G. R. METAL: fast and efficient meta-analysis of genomewide association scans. Bioinformatics 26, 2190–1 (2010).

43. Evangelou, E. & Ioannidis, J. P. A. Meta-analysis methods for genome-wide association studies and beyond. Nat. Rev. Genet. 14, 379–89 (2013).

44. Yang, J., Lee, S. H., Goddard, M. E. & Visscher, P. M. GCTA: a tool for genome-wide complex trait analysis. Am. J. Hum. Genet. 88, 76–82 (2011).

45. Yang, J. et al. Conditional and joint multiple-SNP analysis of GWAS summary statistics identifies additional variants influencing complex traits. Nature Genetics vol. 44 369–375 (2012).

46. Manichaikul, A. et al. Robust relationship inference in genome-wide association studies. Bioinformatics 26, 2867–73 (2010).

47. Pers, T. H. et al. Biological interpretation of genome-wide association studies using predicted gene functions. Nat. Commun. 6, 5890 (2015).

48. Yang, H., Robinson, P. N. & Wang, K. Phenolyzer: phenotype-based prioritization of candidate genes for human diseases. Nat. Methods 12, 841–3 (2015).

49. Zhu, Z. et al. Integration of summary data from GWAS and eQTL studies predicts complex trait gene targets. Nat. Genet. 48, 481–487 (2016).

50. Võsa, U. et al. Unraveling the polygenic architecture of complex traits using blood eQTL meta-analysis. bioRxiv 447367 (2018) doi:10.1101/447367.

51. Qi, T. et al. Identifying gene targets for brain-related traits using transcriptomic and methylomic data from blood. Nat. Commun. 9, 2282 (2018).

52. Vilhjálmsson, B. J. et al. Modeling Linkage Disequilibrium Increases Accuracy of Polygenic Risk Scores. Am. J. Hum. Genet. 97, 576–92 (2015).

53. Bitarello, B. D. et al. Signatures of Long-Term Balancing Selection in Human Genomes. Genome Biol. Evol. 10, 939–955 (2018).

54. 1000 Genomes Project Consortium et al. A global reference for human genetic variation. Nature 526, 68–74 (2015).

