## Supplementary Figures and Text for "Genome-wide analysis identifies genetic effects on reproductive success and ongoing natural selection at the *FADS* locus"

**Figure S1 | Association of the 43 identified loci on NEB and childlessness.** The direction of effect has been reversed for childless to show a risk increasing odds-ratio for easier interpretation of effect magnitude.

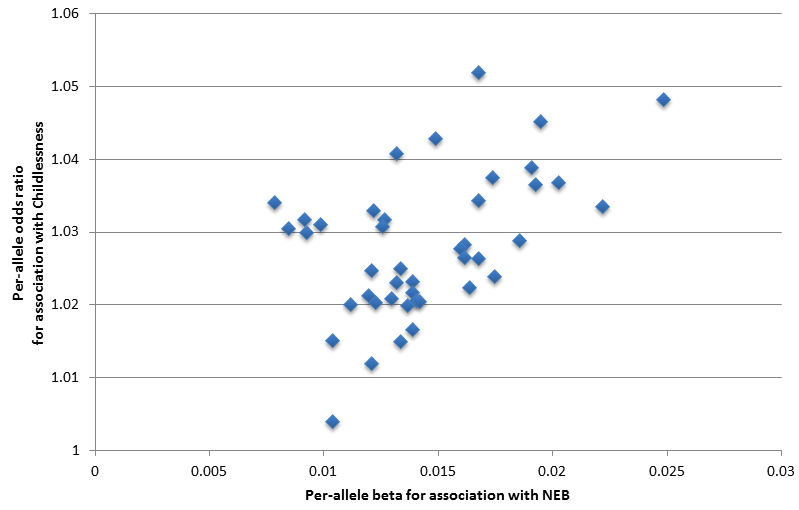

**Figure S2 | Candidate genes in the 43 NEB and childlessness associated loci.** Information for 54 genes prioritized in loci identified by GWAS for number of children ever born (NEB) and/or childlessness (CLN) that are located within 1 million bp of lead SNPs. Blue and orange indicate transitions from one locus to the next. From left to right, panels indicate: 1) if the locus was identified for NEB and/or CLN; 2) which bioinformatic approaches highlighted the gene as a candidate; 3) the cell types in brain, glands, female reproductive organs, and male reproductive organs in which the genes are expressed at a low, moderate or high level (small, medium and large circles) based on data from the Human Protein Atlas; 4) gene functions as extracted from Entrez, Uniprot and GeneCards; 5) which phenotypes were observed in mutant
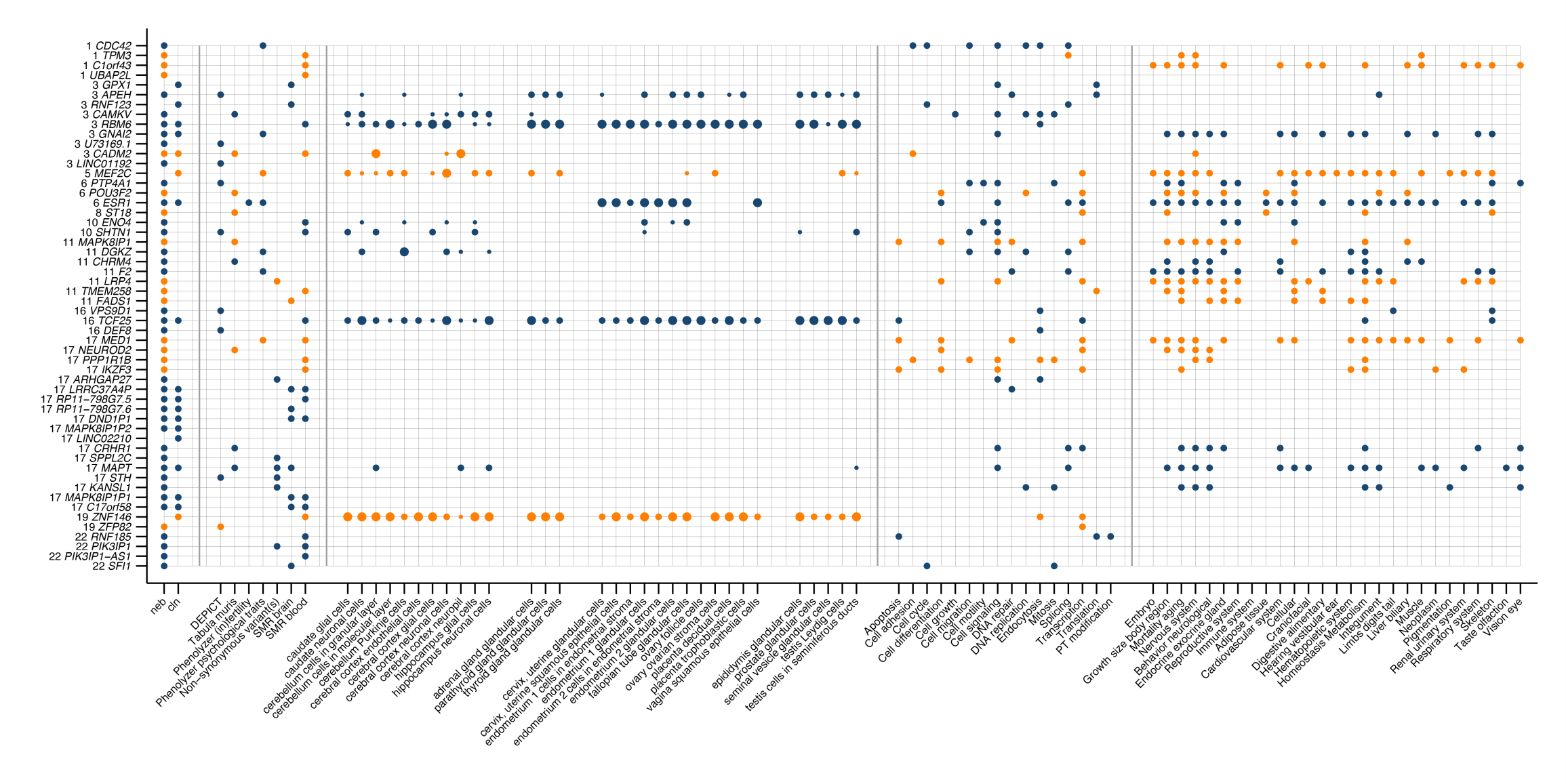

**Figure S3 | Change in polygenic risk score (PGS) for NEB by birth year in the HRS study**. Polygenic risk score scaled to have a mean of 0 and standard deviation of 1.

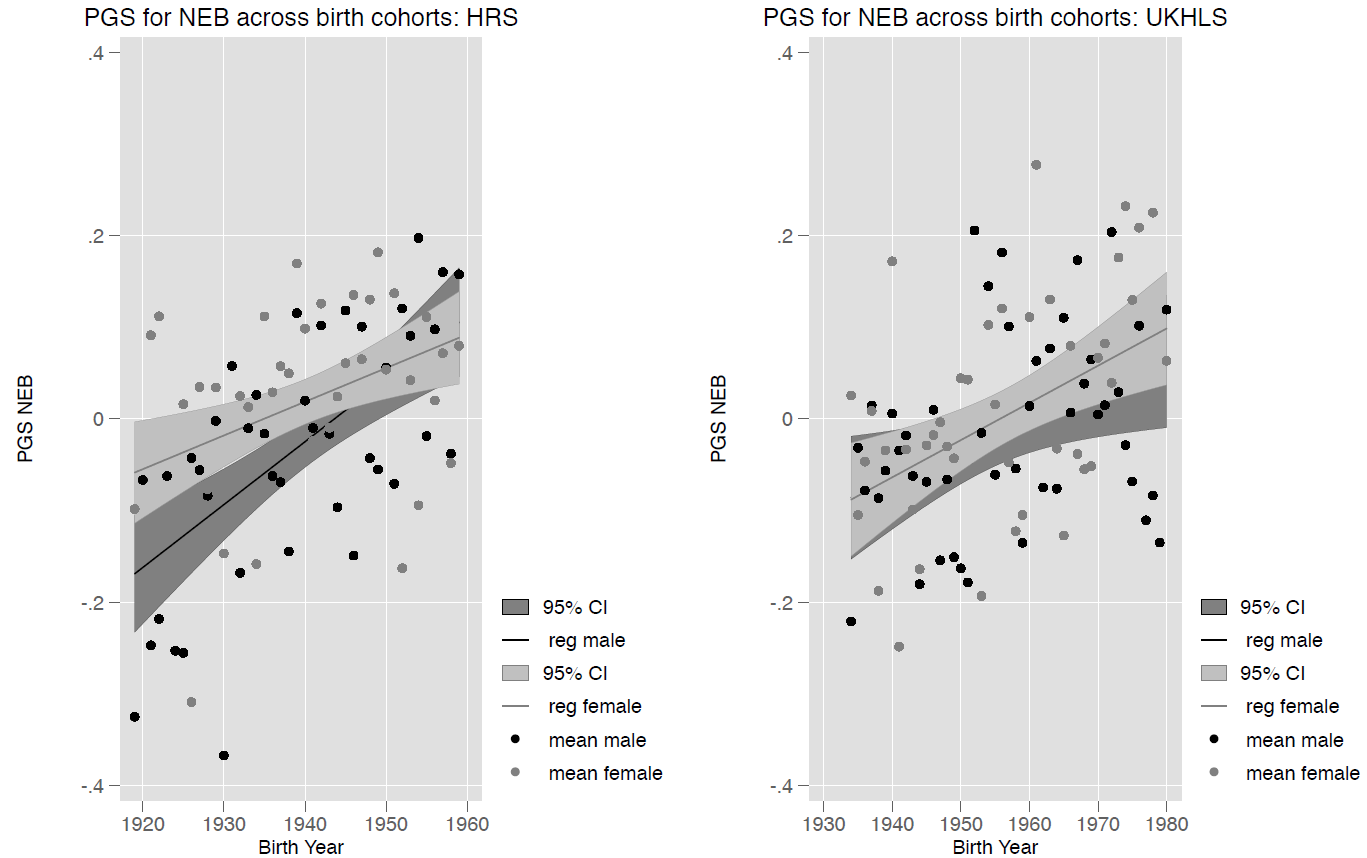

**Human Reproductive Behaviour Consortium - Author information**

Evelina T. Akimova^1^, Sven Bergmann^2,3,4^, Lawrence F. Bielak^5^, Dorret I. Boomsma^6^, Kristina Bosak^7^, Marco Brumat^8^, Julie E. Buring^9,10^, David Cesarini^11,12.13^, Daniel I. Chasman^9,10^, Jorge E. Chavarro^14,15,16^, Massimiliano Cocca^17^, Maria Pina Concas^17^, George Davey-Smith^18^, Gail Davies^19^, Ian J. Deary^19^, Tõnu Esko^20,21^, Jessica D. Faul^22^, Oscar Franco^23^, Andrea Ganna^24,25^, Audrey J. Gaskins^15,16,26^, Andrea Gelemanović^27^, Eco J.C. de Geus^6^, Christian Gieger^28^, Giorgia Girotto^8,17^, Bamini Gopinath^29^, Hans Jörgen Grabe^30^, Erica P. Gunderson^31^, Caroline Hayward^32^, Chunyan He^33,34^, Diana van Heemst^35^, W. David Hill^19^, Georg Homuth^36^, Jouke Jan Hottenga^6^, Hongyang Huang^14^, Elina Hyppӧnen^37,38^, M. Arfan Ikram^23^, Rick Jansen^39^, Magnus Johannesson^40^, Zoha Kamali^41^, Sharon L.R. Kardia^5^, Maryam Kavousi^23^, Annette Kifley^29^, Tuomo Kiiskinen^24,42^, Peter Kraft^14,43^, Brigitte Kühnel^28^, Claudia Langenberg^44^, Gerald Liew^29^, Lifelines Cohort Study^45,46^, Penelope A. Lind^47^, Jian’an Luan^44^, Reedik Mägi^20^, Patrik K.E. Magnusson^48^, Anubha Mahajan^49,50^, Nicholas G. Martin^51^, Hamdi Mbarek^6,52^, Mark I. McCarthy^49,50^, George McMahon^53^, Sarah E. Medland^47^, Thomas Meitinger^54^, Andres Metspalu^20,55^, Evelin Mihailov^20^, Lili Milani^20^, Stacey A. Missmer^14,56,57^, Paul Mitchell^29^, Stine Møllegaard^58^, Dennis O. Mook-Kanamori^59,60^, Anna Morgan^17^, Peter J. van der Most^45^, Renée de Mutsert^59^, Matthias Nauck^61^, Ilja M. Nolte^45^, Raymond Noordam^35^, Brenda W.J.H. Penninx^62^, Annette Peters^63^, Patricia A. Peyser^5^, Ozren Polašek^27,64^, Chris Power^65^, Ajka Pribisalić^27^, Paul Redmond^19^, Janet W. Rich-Edwards^14,16,66^, Paul M. Ridker^9,10^, Cornelius A. Rietveld^67,68^, Susan M. Ring^18^, Lynda M. Rose^9^, Rico Rueedi^2,3^, Jennifer A. Smith^5^, Kári Stefánsson^69^, Doris Stöckl^63^, Konstantin Strauch^70,71,72^, Morris A. Swertz^46^, Alexander Teumer^73^, Gudmar Thorleifsson^69^, Unnur Thorsteinsdottir^69^, A. Roy Thurik^67,68,74^, Nicholas J. Timpson^18^, Constance Turman^14^, André G. Uitterlinden^67,75^, Melanie Waldenberger^28,63^, Nicholas J. Wareham^44^, David R. Weir^22^, Gonneke Willemsen^6^, Wei Zhao^5^, and Jing Hau Zhao^44^

Author list is ordered alphabetically.

^1^ Leverhulme Centre for Demographic Science, Department of Sociology, St. Antony’s College, University of Oxford, Oxford, United Kingdom

^2^ Department of Computational Biology, University of Lausanne, Lausanne, Switzerland

^3^ Swiss Institute of Bioinformatics, Lausanne, Switzerland

^4^ Department of Integrative Biomedical Sciences, University of Cape Town, Cape Town, South Africa

^5^ Department of Epidemiology, University of Michigan, Ann Arbor, MI, United States of America

^6^ Department of Biological Psychology, Amsterdam Public Health Research Institute, Vrije Universiteit Amsterdam, Amsterdam, The Netherlands

^7^ Psychiatric hospital “Sveti Ivan”, Zagreb, Croatia

^8^ Department of Medical, Surgical and Health Sciences, University of Trieste, Trieste, Italy

^9^ Brigham and Women’s Hospital, Boston, MA, United States of America

^10^ Harvard Medical School, Boston, MA, United States of America

^11^ Department of Economics, New York University, New York, NY, United States of America

^12^ Research Institute for Industrial Economics, Stockholm, Sweden

^13^ National Bureau of Economic Research, Cambridge, MA, United States of America

^14^ Department of Epidemiology, Harvard T.H. Chan School of Public Health, Boston, MA, United States of America

^15^ Department of Nutrition, Harvard T.H. Chan School of Public Health, Boston, MA, United States of America

^16^ Channing Division of Network Medicine, Brigham and Women’s Hospital and Harvard Medical School, Boston, MA, United States of America

^17^ Institute for Maternal and Child Health IRCCS “Burlo Garofolo”, Trieste, Italy

^18^ MRC Integrative Epidemiology Unit, University of Bristol, Bristol, United Kingdom

^19^ Lothian Birth Cohorts, Department of Psychology, University of Edinburgh, Edinburgh, United Kingdom

^20^ Estonian Genome Center, University of Tartu, Tartu, Estonia

^21^ Broad Institute of the Massachusetts Institute of Technology and Harvard University, Cambridge, MA, United States of America

^22^ Survey Research Center, Institute for Social Research, University of Michigan, Ann Arbor, MI, United States of America

^23^ Department of Epidemiology, Erasmus Medical Center, Rotterdam, The Netherlands

^24^ Institute for Molecular Medicine Finland (FIMM), HiLIFE, University of Helsinki, Helsinki, Finland

^25^ Analytic and Translational Genetics Unit, Center for Genomic Medicine, Massachusetts General Hospital, Boston, MA, United States of America

^26^ Department of Epidemiology, Rollins School of Public Health, Emory University, Atlanta, GA, United States of America

^27^ University of Split School of Medicine, Split, Croatia

^28^ Research Unit of Molecular Epidemiology, Helmholtz Zentrum München, German Research Center for Environmental Health, Neuherberg, Germany

^29^ Centre for Vision Research, Westmead Institute for Medical Research and Department of Ophthalmology, University of Sydney, Sydney, Australia

^30^ Department of Psychiatry and Psychotherapy, University Medicine Greifswald, Greifswald, Germany

^31^ Division of Research, Kaiser Permanente Northern California, Oakland, CA, United States of America

^32^ Medical Research Council Human Genetics Unit, Institute of Genetics and Molecular Medicine, University of Edinburgh, Edinburgh, United Kingdom

^33^ University of Kentucky Markey Cancer Center, Lexington, KY, United States of America

^34^ Department of Internal Medicine, Division of Medical Oncology, University of Kentucky College of Medicine, Lexington, KY, United States of America

^35^ Department of Internal Medicine, Section of Gerontology and Geriatrics, Leiden University Medical Center, Leiden, The Netherlands

^36^ Interfaculty Institute for Genetics and Functional Genomics, University of Greifswald, Greifswald, Germany

^37^ Australian Centre for Precision Health, University of South Australia Cancer Research Institute, Adelaide, Australia

^38^ South Australian Health and Medical Research Institute, Adelaide, Australia

^39^ Department of Psychiatry, Amsterdam Public Health and Amsterdam Neuroscience, Amsterdam UMC, Vrije Universiteit, Amsterdam, The Netherlands

^40^ Department of Economics, Stockholm School of Economics, Stockholm, Sweden

^41^ Department of Bioinformatics, Isfahan University of Medical Sciences, Isfahan, Iran

^42^ National Institute for Health and Welfare, Helsinki, Finland

^43^ Department of Biostatistics, Harvard T.H. Chan School of Public Health, Boston, MA, United States of America

^44^ MRC Epidemiology Unit, Institute of Metabolic Science, Cambridge Biomedical Campus, University of Cambridge School of Clinical Medicine, Cambridge, United Kingdom

^45^ Department of Epidemiology, University of Groningen, University Medical Center Groningen, Groningen, The Netherlands

^46^ Department of Genetics, University of Groningen, University Medical Center Groningen, Groningen, The Netherlands

^47^ Psychiatric Genetics, QIMR Berghofer Medical Research Institute, Herston Brisbane, Queensland, Australia

^48^ Department of Medical Epidemiology and Biostatistics, Karolinska Institutet, Stockholm, Sweden

^49^ Wellcome Centre for Human Genetics, University of Oxford, Oxford, United Kingdom

^50^ Oxford Centre for Diabetes, Endocrinology and Metabolism, Radcliffe Department of Medicine, University of Oxford, Oxford, United Kingdom

^51^ Genetic Epidemiology, QIMR Berghofer Medical Research Institute, Herston Brisbane, Queensland, Australia

^52^ Qatar Genome Programme, Qatar Foundation, Doha, Qatar

^53^ School of Social and Community Medicine University of Bristol, Bristol, United Kingdom

^54^ Institute of Human Genetics, Helmholtz Zentrum München, German Research Center for Environmental Health, Neuherberg, Germany

^55^ Institute of Molecular and Cell Biology, University of Tartu, Tartu, Estonia

^56^ Division of Adolescent and Young Adult Medicine, Department of Medicine, Boston Children’s Hospital and Harvard Medical School, Boston, MA, United States of America

^57^ Department of Obstetrics, Gynecology, and Reproductive Biology, College of Human Medicine, Michigan State University, Grand Rapids, MI, United States of America

^58^ Department of Sociology, University of Copenhagen, Copenhagen, Denmark

^59^ Department of Clinical Epidemiology, Leiden University Medical Center, Leiden, The Netherlands

^60^ Department of Public Health and Primary Care, Leiden University Medical Center, Leiden, The Netherlands

^61^ Institute of Clinical Chemistry and Laboratory Medicine, University Medicine Greifswald, Greifswald, Germany

^62^ Department of Psychiatry, EMGO Institute for Health and Care Research and Neuroscience Campus Amsterdam, VU University Medical Center/GGZ inGeest, Amsterdam, The Netherlands

^63^ Institute of Epidemiology, Helmholtz Zentrum München, German Research Center for Environmental Health, Neuherberg, Germany

^64^ Gen Info, LLC, Zagreb, Croatia

^65^ Population, Policy and Practice Research and Teaching Department, UCL Great Ormond Street Institute of Child Health, London, United Kingdom

^66^ Division of Women’s Health, Department of Medicine, Brigham and Women’s Hospital and Harvard Medical School, Boston, MA, United States of America

^67^ Erasmus University Rotterdam Institute for Behavior and Biology, Rotterdam, The Netherlands

^68^ Department of Applied Economics, Erasmus School of Economics, Rotterdam, The Netherlands

^69^ deCODE Genetics/Amgen Inc., Reykjavik, Iceland

^70^ Institute of Medical Biostatistics, Epidemiology and Informatics (IMBEI), University Medical Center, Johannes Gutenberg University, Mainz, Germany

^71^ Institute of Genetic Epidemiology, Helmholtz Zentrum München, German Research Center for Environmental Health, Neuherberg, Germany

^72^ Chair of Genetic Epidemiology, IBE, Faculty of Medicine, LMU Munich, Germany

^73^ Institute for Community Medicine, University Medicine Greifswald, Greifswald, Germany

^74^ Montpellier Business School, Montpellier, France

^75^ Department of Internal Medicine, Erasmus University Medical Center, Rotterdam, The Netherlands

**eQTLGen Consortium**

Mawussé Agbessi^1^, Habibul Ahsan^2^, Isabel Alves^1^, Anand Andiappan^3^, Wibowo Arindrarto^4^, Philip Awadalla^1^, Alexis Battle^5,6^, Frank Beutner^7^, Marc Jan Bonder^8,9,10^, Dorret Boomsma^11^, Mark Christiansen^12^, Annique Claringbould^8^, Patrick Deelen^8,13^, Tõnu Esko^14^, Marie-Julie Favé^1^, Lude Franke^8^, Timothy Frayling^15^, Sina A. Gharib^12,16^, Gregory Gibson^17^, Bastiaan T. Heijmans^4^, Gibran Hemani^18^, Rick Jansen^19^, Mika Kähönen^20^, Anette Kalnapenkis^14^, Silva Kasela^14^, Johannes Kettunen^21^, Yungil Kim^6,22^, Holger Kirsten^23^, Peter Kovacs^24^, Knut Krohn^25^, Jaanika Kronberg-Guzman^14^, Viktorija Kukushkina^14^, Zoltan Kutalik^26^, Bernett Lee^3^, Terho Lehtimäki^27^, Markus Loeffler^23^, Urko M. Marigorta^17^, Hailang Mei^4^, Lili Milani^14^, Grant W. Montgomery^28^, Martina Müller-Nurasyid^29,30,31^, Matthias Nauck^32^, Michel Nivard^11^, Brenda Penninx^19^, Markus Perola^33^, Natalia Pervjakova^14^, Brandon L. Pierce^2^, Joseph Powell^34^, Holger Prokisch^35,36^, Bruce M. Psaty^12,37,38^, Olli T. Raitakari^39^, Samuli Ripatti^40^, Olaf Rotzschke^3^, Sina Rüeger^26^, Ashis Saha^6^, Markus Scholz^23^, Katharina Schramm^31,32^, Ilkka Seppälä^27^, Eline P. Slagboom^4^, Coen D.A. Stehouwer^41^, Michael Stumvoll^42^, Patrick Sullivan^43^, Peter A.C. ‘t Hoen^44^, Alexander Teumer^45^, Joachim Thiery^46^, Lin Tong^2^, Anke Tönjes^42^, Jenny van Dongen^11^, Maarten van Iterson^4^, Joyce van Meurs^47^, Jan H. Veldink^48^, Joost Verlouw^47^, Peter M. Visscher^28^, Uwe Völker^49^, Urmo Võsa^8,14^, Harm-Jan Westra^8^, Cisca Wijmenga^8^, Hanieh Yaghootkar^15^, Jian Yang^28,50^, Biao Zeng^17^, & Futao Zhang^28^

*Author list is ordered alphabetically.*

^1^ Computational Biology, Ontario Institute for Cancer Research, Toronto, Canada

^2^ Department of Public Health Sciences, University of Chicago, Chicago, United States of America

^3^ Singapore Immunology Network, Agency for Science, Technology and Research, Singapore, Singapore

^4^ Department of Biomedical Data Sciences, Leiden University Medical Center, Leiden, The Netherlands

^5^ Department of Biomedical Engineering, Johns Hopkins University, Baltimore, United States of America

^6^ Department of Computer Science, Johns Hopkins University, Baltimore, United States of America

^7^ Heart Center Leipzig, Universität Leipzig, Leipzig, Germany

^8^ Department of Genetics, University Medical Centre Groningen, Groningen, The Netherlands

^9^ European Molecular Biology Laboratory, European Bioinformatics Institute, Wellcome Genome Campus, Hinxton, United Kingdom

^10^ Genome Biology Unit, European Molecular Biology Laboratory, Heidelberg, Germany

^11^ Department of Biological Psychology, Vrije Universiteit Amsterdam, Amsterdam, The Netherlands

^12^ Cardiovascular Health Research Unit, University of Washington, Seattle, United States of America

^13^ Genomics Coordination Center, University Medical Centre Groningen, Groningen, The Netherlands

^14^ Estonian Genome Center, Institute of Genomics, University of Tartu, Tartu 51010, Estonia

^15^ Exeter Medical School, University of Exeter, Exeter, United Kingdom

^16^ Department of Medicine, University of Washington, Seattle, United States of America

^17^ School of Biological Sciences, Georgia Tech, Atlanta, United States of America

^18^ MRC Integrative Epidemiology Unit, University of Bristol, Bristol, United Kingdom

^19^ Department of Psychiatry and Amsterdam Neuroscience, Amsterdam UMC, Vrije Universiteit, Amsterdam, the Netherlands

^20^ Department of Clinical Physiology, Tampere University Hospital and Faculty of Medicine and Health Technology, Tampere University, Tampere, Finland

^21^ Centre for Life Course Health Research, University of Oulu, Oulu, Finland

^22^ Genetics and Genomic Science Department, Icahn School of Medicine at Mount Sinai, New York, United States of America

^23^ Institut für Medizinische Informatik, Statistik und Epidemiologie, LIFE – Leipzig Research Center for Civilization Diseases, Universität Leipzig, Leipzig, Germany

^24^ IFB Adiposity Diseases, Universität Leipzig, Leipzig, Germany

^25^ Interdisciplinary Center for Clinical Research, Faculty of Medicine, Universität Leipzig, Leipzig, Germany

^26^ Institute of Social and Preventive Medicine, Lausanne University Hospital, Lausanne, Switzerland

^27^ Department of Clinical Chemistry, Fimlab Laboratories and Finnish Cardiovascular Research Center-Tampere, Faculty of Medicine and Health Technology, Tampere University, Tampere, Finland

^28^ Institute for Molecular Bioscience, University of Queensland, Brisbane, Australia

^29^ Institute of Genetic Epidemiology, Helmholtz Zentrum München - German Research Center for Environmental Health, Neuherberg, Germany

^30^ Department of Medicine I, University Hospital Munich, Ludwig Maximilian’s University, Munich, Germany

^31^ DZHK (German Centre for Cardiovascular Research), partner site Munich Heart Alliance, Munich, Germany

^32^ Institute of Clinical Chemistry and Laboratory Medicine, University Medicine Greifswald, Greifswald, Germany

^33^ National Institute for Health and Welfare, University of Helsinki, Helsinki, Finland

^34^ Garvan Institute of Medical Research, Garvan-Weizmann Centre for Cellular Genomics, Sydney, Australia

^35^ Institute of Human Genetics, Helmholtz Zentrum München, Neuherberg, Germany

^36^ Institute of Human Genetics, Technical University Munich, Munich, Germany

^37^ Departments of Epidemiology, Medicine, and Health Services, University of Washington, Seattle, United States of America

^38^ Kaiser Permanente Washington Health Research Institute, Seattle, WA, United States of America

^39^ Centre for Population Health Research, Department of Clinical Physiology and Nuclear Medicine, Turku University Hospital and University of Turku, Turku, Finland

^40^ Statistical and Translational Genetics, University of Helsinki, Helsinki, Finland

^41^ Department of Internal Medicine, Maastricht University Medical Centre, Maastricht, The Netherlands

^42^ Department of Medicine, Universität Leipzig, Leipzig, Germany

^43^ Department of Medical Epidemiology and Biostatistics, Karolinska Institutet, Stockholm, Sweden

^44^ Center for Molecular and Biomolecular Informatics, Radboud Institute for Molecular Life Sciences, Radboud University Medical Center Nijmegen, Nijmegen, The Netherlands

^45^ Institute for Community Medicine, University Medicine Greifswald, Greifswald, Germany

^46^ Institute for Laboratory Medicine, LIFE – Leipzig Research Center for Civilization Diseases, Universität Leipzig, Leipzig, Germany

^47^ Department of Internal Medicine, Erasmus Medical Centre, Rotterdam, The Netherlands

^48^ Department of Neurology, University Medical Center Utrecht, Utrecht, The Netherlandsy

^49^ Interfaculty Institute for Genetics and Functional Genomics, University Medicine Greifswald, Greifswald, Germany

^50^ Institute for Advanced Research, Wenzhou Medical University, Wenzhou, Zhejiang 325027, China

**Acknowledgements**

The research leading to these results has received funding from the following awards to PI M.C. Mills, European Research Council (ERC) Consolidator Grant SOCIOGENOME (615603, [www.sociogenome.org](http://www.sociogenome.org)), Economic & Social Research Council (ESRC) UK, National Centre for Research Methods (NCRM) grant SOCGEN, Wellcome Trust ISSF and John Fell Grant and a large Centre grant from the Leverhulme Trust for the Leverhulme Centre for Demographic Science.

**1958BC-T1DGC and 1958BC-WTCCC2**

This work made use of data and samples generated by the 1958 Birth Cohort (NCDS), which is managed by the Centre for Longitudinal Studies at the UCL Institute of Education, funded by the Economic and Social Research Council (grant number ES/M001660/1). Data governance was provided by the METADAC data access committee, funded by ESRC, Wellcome, and MRC. (2015-2018: Grant Number MR/N01104X/1 2018-2020: Grant Number ES/S008349/1). Access to these resources was enabled via the Wellcome Trust & MRC: 58FORWARDS grant [108439/Z/15/Z] (The 1958 Birth Cohort: Fostering new Opportunities for Research via Wider Access to Reliable Data and Samples). Before 2015 biomedical resources were maintained under the Wellcome Trust and Medical Research Council 58READIE Project (grant numbers WT095219MA and G1001799). Genotyping was undertaken as part of the Wellcome Trust Case-Control Consortium (WTCCC) under Wellcome Trust award 076113, and a full list of the investigators who contributed to the generation of the data is available at www.wtccc.org.uk. This research used resources provided by the Type 1 Diabetes Genetics Consortium, a collaborative clinical study sponsored by the National Institute of Diabetes and Digestive and Kidney Diseases (NIDDK), National Institute of Allergy and Infectious Diseases, National Human Genome Research Institute, National Institute of Child Health and Human Development, and Juvenile Diabetes Research Foundation International (JDRF) and supported by U01 DK062418. The 1958 birth cohort data can be accessed via the UK Data Service (<http://ukdataservice.ac.uk/>). Funding: Niddk. U01-DK105535; Wellcome: 090532, 098381, 106130, 203141, 212259; MMcC was a Wellcome Investigator and an NIHR Senior Investigator

**ALSPAC: Avon Longitudinal Study of Parent and Children**

We are extremely grateful to all the families who took part in this study, the midwives for their help in recruiting them, and the whole ALSPAC team, which includes interviewers, computer and laboratory technicians, clerical workers, research scientists, volunteers, managers, receptionists and nurses. The UK Medical Research Council and Wellcome (Grant ref: 217065/Z/19/Z) and the University of Bristol provide core support for ALSPAC. This publication is the work of the authors and will serve as guarantors for the contents of this paper. A comprehensive list of grants funding is available on the ALSPAC website (<http://www.bristol.ac.uk/alspac/external/documents/grant-acknowledgements.pdf>). GDS works in the Medical Research Council Integrative Epidemiology Unit at the University of Bristol (MC_UU_00011/1)

**BMES: The Blue Mountains Eye Study**

The Blue Mountains Eye Study (BMES) was supported by the Australian National Health & Medical Research Council (NHMRC), Canberra Australia (NHMRC project grant IDs 974159, 211069, 302068, and Centre for Clinical Research Excellence in Translational Clinical Research in Eye Diseases, CCRE in TCR-Eye, grant ID 529923). The BMES GWAS and genotyping costs was supported by Australian NHMRC, Canberra Australia (NHMRC project grant IDs 512423, 475604 and 529912), and the Wellcome Trust, UK as part of Wellcome Trust Case Control Consortium 2 (A Viswanathan, P McGuffin, P Mitchell, F Topouzis, P Foster, grant IDs 085475/B/08/Z and 085475/08/Z).

**coLaus: Cohorte Lausannoise**

The CoLaus study was and is supported by research grants from GlaxoSmithKline (GSK), the Faculty of Biology and Medicine of Lausanne, and the Swiss National Science Foundation (grants 3200B0-105993, 3200B0- 118308, 33CSCO-122661, and 33CS30-139468). We thank all participants, involved physicians and study nurses to the CoLaus cohort.

**CROATIA cohorts**

We would like to acknowledge the invaluable contributions of the recruitment team in Korcula, the administrative teams in Croatia and Edinburgh and the people of Korcula. The CROATIA-Korcula study was funded by grants from the Medical Research Council (UK), European Commission Framework 6 project EUROSPAN (Contract No. LSHG-CT-2006-018947), FP7 project BBMRI-LPC (grant 313010), Ministry of Science, Education and Sports of the Republic of Croatia (grant 108-1080315-0302), the Croatian Science Foundation (grant 8875), the Croatian National Centre of Research Excellence in Personalized Healthcare grant (number KK.01.1.1.01.0010) and the Centre of Competence in Molecular Diagnostics (KK.01.2.2.03.0006). External researchers who wish to obtain access to CROATIA-Korcula’s data or EA2 results may contact Ozren Polasek,.

**deCODE**

We thank the study subjects for their valuable participation. All deCODE collaborators in this study are employees of deCODE Genetics/Amgen, Inc. External researchers who wish to obtain access to data or EA2 results may contact Gudmar Thorleifsson.

**EGCUT: Estonian Genome Center, University of Tartu**

EGCUT received funding from the Estonian Research Council Grant IUT20-60 and PUT1660, EU H2020 grant 692145, and European Union through the European Regional Development Fund (Project No. 2014-2020.4.01.15-0012) GENTRANSMED. For more information, please contact Tõnu Esko.

**EPIC Norfolk: The European Prospective Investigation in Cancer and Nutrition Norfolk study**

The authors would like to acknowledge the contribution of the staff and participants of the EPIC-Norfolk Study. EPIC-Norfolk is supported by the Medical Research Council (programme grants G0401527, G1000143) and Cancer Research UK (programme grant C864/A8257). This work was supported by the Medical Research Council (Unit Programme numbers MC_UU_12015/1 and MC_UU_12015/2). For inquiries about access to this data, please contact Ken Ong.

**FinnGen**

The FinnGen project is funded by two grants from Business Finland (HUS 4685/31/2016 and UH 4386/31/2016) and nine industry partners (AbbVie, AstraZeneca, Biogen, Celgene, Genentech, GSK, MSD, Pfizer and Sanofi). Following biobanks are acknowledged for collecting the FinnGen project samples: Auria Biobank (https://www.auria.fi/biopankki/en), THL Biobank (https://thl.fi/fi/web/thl-biopankki), Helsinki Biobank (https://www.terveyskyla.fi/helsinginbiopankki/en), Northern Finland Biobank Borealis (https://www.ppshp.fi/Tutkimus-ja-opetus/Biopankki), Finnish Clinical Biobank Tampere (https://www.tays.fi/en-US/Research_and_development/Finnish_Clinical_Biobank_Tampere), Biobank of Eastern Finland (https://ita-suomenbiopankki.fi/), Central Finland Biobank (https://www.ksshp.fi/fi-FI/Potilaalle/Biopankki), Finnish Red Cross Blood Service Biobank (https://www.bloodservice.fi/Research%20Projects/biobanking).

**GENOA: Genetic Epidemiology Network of Arteriopathy**

Support for GENOA was provided by the National Heart, Lung and Blood Institute (HL119443, HL118305, HL054464, HL054457, HL054481, HL071917 and HL87660) of the National Institutes of Health. Genotyping was performed at the Mayo Clinic (Stephen T. Turner, MD, Mariza de Andrade PhD, Julie Cunningham, PhD). We thank Eric Boerwinkle, PhD and Megan L. Grove from the Human Genetics Center and Institute of Molecular Medicine and Division of Epidemiology, University of Texas Health Science Center, Houston, Texas, USA for their help with genotyping. We would also like to thank the families that participated in the GENOA study.

**HRS: Health and Retirement Study**

HRS is supported by the National Institute on Aging (NIA U01AG009740). The genotyping was funded separately by the National Institute on Aging (RC2 AG036495, RC4 AG039029). Our genotyping was conducted by the NIH Center for Inherited Disease Research (CIDR) at Johns Hopkins University. Genotyping quality control and final preparation of the data were performed by the Genetics Coordinating Center at the University of Washington.

**INGI-FVG: Friuli Venezia Giulia Genetic Park**

We would like to thank the people of the Friuli Venezia Giulia Region for the everlasting support. The research was supported by Italian Ministry of Health - RC 35/17.

**InterAct-GWAS and InterAct-Exome**

We thank all EPIC participants and staff for their contribution to the study. We thank Nicola Kerrison (MRC Epidemiology Unit, Cambridge) for managing the data for the InterAct Project. Funding for the InterAct project was provided by the EU FP6 programme (grant number LSHM_CT_2006_037197).

**KORA F3 and F4**

The KORA study was initiated and financed by the Helmholtz Zentrum München – German Research Center for Environmental Health, which is funded by the German Federal Ministry of Education and Research (BMBF) and by the State of Bavaria. Furthermore, KORA research was supported within the Munich Center of Health Sciences (MC-Health), Ludwig-Maximilians-Universität, as part of LMUinnovativ. The funders had no role in study design, data collection and analysis, decision to publish, or preparation of the manuscript. We thank all the study participants, all members of staff of the Institute of Epidemiology II and the field staff in Augsburg who planned and conducted the study.

**LBC1921 and LBC1936: The Lothian Birth Cohorts**

We thank the cohort participants and team members who contributed to these studies. Phenotype collection in the Lothian Birth Cohort 1921 was supported by the UK’s Biotechnology and Biological Sciences Research Council (BBSRC), The Royal Society, and The Chief Scientist Office of the Scottish Government. Phenotype collection in the Lothian Birth Cohort 1936 was supported by Age UK (The Disconnected Mind project). Genotyping of the cohorts was funded by the BBSRC. The work was undertaken by The University of Edinburgh Centre for Cognitive Ageing and Cognitive Epidemiology, part of the cross council Lifelong Health and Wellbeing Initiative (MR/K026992/1). Funding from the BBSRC and Medical Research Council (MRC) is gratefully acknowledged. WDH is supported from a grant from Age UK (The Disconnected Mind Project).

**Lifelines Cohort Study**

We wish to acknowledge the services of the Lifelines Cohort Study, the contributing research centers delivering data to Lifelines, and all the study participants. The Lifelines Cohort Study, and generation and management of GWAS genotype data for the Lifelines Cohort Study is supported by the Netherlands Organization of Scientific Research NWO (grant 175.010.2007.006), the Economic Structure Enhancing Fund (FES) of the Dutch government, the Ministry of Economic Affairs, the Ministry of Education, Culture and Science, the Ministry for Health, Welfare and Sports, the Northern Netherlands Collaboration of Provinces (SNN), the Province of Groningen, University Medical Center Groningen, the University of Groningen, Dutch Kidney Foundation and Dutch Diabetes Research Foundation. We thank Behrooz Alizadeh, Annemieke Boesjes, Marcel Bruinenberg, Noortje Festen, Pim van der Harst, Ilja Nolte, Lude Franke, Mitra Valimohammadi for their help in creating the GWAS database, and Rob Bieringa, Joost Keers, René Oostergo, Rosalie Visser, Judith Vonk for their work related to data-collection and validation. The authors are grateful to the study participants, the staff from the LifeLines Cohort Study and the contributing research centers delivering data to LifeLines and the participating general practitioners and pharmacists.

**NEO: Netherlands Epidemiology of Obesity**

The authors of the NEO study thank all individuals who participated in the Netherlands Epidemiology in Obesity study, all participating general practitioners for inviting eligible participants and all research nurses for collection of the data. We thank the NEO study group, Pat van Beelen, Petra Noordijk and Ingeborg de Jonge for the coordination, lab and data management of the NEO study. The genotyping in the NEO study was supported by the Centre National de Génotypage (Paris, France), headed by Jean-Francois Deleuze. The NEO study is supported by the participating Departments, the Division and the Board of Directors of the Leiden University Medical Center, and by the Leiden University, Research Profile Area Vascular and Regenerative Medicine. Dennis Mook-Kanamori is supported by Dutch Science Organization (ZonMW-VENI Grant 916.14.023).

**NESDA: The Netherlands Study of Depression and Anxiety**

Funding was obtained from the Netherlands Organization for Scientific Research (Geestkracht program grant 10-000-1002); the Center for Medical Systems Biology (CSMB, NOW Genomics), Biobanking and Biomolecular Resources Research Infrastructure (BBMRI-NL), VU University’s Institutes for Health and Care Research (EMGO+) and Neuroscience Campus Amsterdam, University Medical Center Groningen, Leiden University Medical Center, National Institutes of Health (NIH, R01D0042157-01A, MH081802, Grand Opportunity grants 1RC2 MH089951 and 1RC2 MH089995). Part of the genotyping and analyses were funded by the Genetic Association Information Network (GAIN) of the Foundation for the National Institutes of Health. Computing was supported by BiG Grid, the Dutch e-Science Grid, which is financially supported by NWO. We would like to thank the Center for Information Technology of the University of Groningen for their support and for providing access to the Peregrine high performance computing cluster.

**NHS: The Nurses’ Health Study**

Supported by grants UM1 CA186107, UM1 CA167552, DK091718, HL071981, HL073168, CA87969, CA49449, CA055075, HL34594, HL088521, U01HG004399, DK080140, 5P30DK46200, U54CA155626, DK58845, U01HG004728-02, EY015473, DK70756 and DK46200 from the National Institutes of Health, with additional support for genotyping from Merck Research Laboratories, North Wales, PA.

**NTR: Netherlands Twin Register**

Funding was obtained from the Netherlands Organization for Scientific Research (NWO) and The Netherlands Organisation for Health Research and Development (ZonMW) grants 904-61-090, 985-10-002, 912-10-020, 904-61-193,480-04-004, 463-06-001, 451-04-034, 400-05-717, Addiction-31160008, 016-115-035, 481-08-011, 400-07-080, 056-32-010, Middelgroot-911-09-032, OCW_NWO Gravity program –024.001.003, NWO-Groot 480-15-001/674, Center for Medical Systems Biology (CSMB, NWO Genomics), NBIC/BioAssist/RK(2008.024), Biobanking and Biomolecular Resources Research Infrastructure (BBMRI –NL, 184.021.007 and 184.033.111), X-Omics 184-034-019; Spinozapremie (NWO- 56-464-14192), KNAW Academy Professor Award (PAH/6635) and University Research Fellow grant (URF) to DIB; Amsterdam Public Health research institute (former EMGO+) , Neuroscience Amsterdam research institute (former NCA) ; the European Community's Fifth and Seventh Framework Program (FP5- LIFE QUALITY-CT-2002-2006, FP7- HEALTH-F4-2007-2013, grant 01254: GenomEUtwin, grant 01413: ENGAGE and grant 602768: ACTION); the European Research Council (ERC Starting 284167, ERC Consolidator 771057, ERC Advanced 230374), Rutgers University Cell and DNA Repository (NIMH U24 MH068457-06), the National Institutes of Health (NIH, R01D0042157-01A1, R01MH58799-03, MH081802, DA018673, R01 DK092127-04, Grand Opportunity grants 1RC2 MH089951, and 1RC2 MH089995); the Avera Institute for Human Genetics, Sioux Falls, South Dakota (USA). Part of the genotyping and analyses were funded by the Genetic Association Information Network (GAIN) of the Foundation for the National Institutes of Health. Computing was supported by NWO through grant 2018/EW/00408559, BiG Grid, the Dutch e-Science Grid and SURFSARA.

**RPGEH: Research Program on Genes, Environment and Health/Genetic Epidemiology Research on Aging (RPGEH/GERA)**

Data used in this study were provided by the Kaiser Permanente Research Program on Genes, Environment, and Health (RPGEH): Genetic Epidemiology Research on Adult Health and Aging (GERA), funded by the National Institutes of Health [RC2 AG036607 (Schaefer and Risch)], the Robert Wood Johnson Foundation, the Wayne and Gladys Valley Foundation, The Ellison Medical Foundation, and the Kaiser Permanente Community Benefits Program. Access to RPGEH data used in this study may be obtained by application via the RPGEH Research portal: https://rpgehportal.kaiser.org. A subset of the GERA cohort consented for public use can be found at NIH/dbGaP: phs000674.v1.p1.

**RS-I (Rotterdam Study Baseline), RS-II (Rotterdam Study Extension of Baseline) and RS-III (Rotterdam Study Young)**

The generation and management of GWAS genotype data for the Rotterdam Study is supported by the Netherlands Organisation of Scientific Research NWO Investments (nr. 175.010.2005.011, 911-03-012). This study is funded by the Research Institute for Diseases in the Elderly (014-93-015; RIDE2), the Netherlands Genomics Initiative (NGI)/Netherlands Organisation for Scientific Research (NWO) project nr. 050-060-810. We thank Pascal Arp, Mila Jhamai, Marijn Verkerk, Lizbeth Herrera and Marjolein Peters for their help in creating the GWAS database, and Karol Estrada and Maksim V. Struchalin for their support in creation and analysis of imputed data. The Rotterdam Study is funded by Erasmus Medical Center and Erasmus University, Rotterdam, Netherlands Organization for the Health Research and Development (ZonMw), the Research Institute for Diseases in the Elderly (RIDE), the Ministry of Education, Culture and Science, the Ministry for Health, Welfare and Sports, the European Commission (DG XII), and the Municipality of Rotterdam. The authors are grateful to the study participants, the staff from the Rotterdam Study and the participating general practitioners and pharmacists. C.A. Rietveld gratefully acknowledges funding from the Netherlands Organization for Scientific Research (NWO Veni grant 016.165.004).

**SHIP: Study of Health in Pomerania**

SHIP is part of the Community Medicine Research net of the University of Greifswald, Germany, which is funded by the Federal Ministry of Education and Research (grants no. 01ZZ9603, 01ZZ0103, and 01ZZ0403), the Ministry of Cultural Affairs as well as the Social Ministry of the Federal State of Mecklenburg-West Pomerania, and the network ‘Greifswald Approach to Individualized Medicine (GANI_MED)’ funded by the Federal Ministry of Education and Research (grant 03IS2061A). Genome-wide data have been supported by the Federal Ministry of Education and Research (grant no. 03ZIK012) and a joint grant from Siemens Healthineers, Erlangen, Germany and the Federal State of Mecklenburg- West Pomerania. The University of Greifswald is a member of the Caché Campus program of the InterSystems GmbH. HJG has received travel grants and speakers honoraria from Fresenius Medical Care, Neuraxpharm, Servier and Janssen Cilag as well as research funding from Fresenius Medical Care.

**STR: Swedish Twin Registry**

The Jan Wallander and Tom Hedelius Foundation (P2015-0001:1), the Ragnar Soderberg Foundation (E9/11, E42/15), The Swedish Research Council (421-2013-1061). STR is financially supported by Karolinska Institutet. Researchers interested in using STR data must obtain approval from a Swedish Ethical Review Board and from the Steering Committee of the Swedish Twin Registry. Researchers using the data are required to follow the terms of an Assistance Agreement containing a number of clauses designed to ensure protection of privacy and compliance with relevant laws. For further information, contact Patrik Magnusson. C.A. Rietveld gratefully acknowledges funding from the Netherlands Organization for Scientific Research (NWO Veni grant 016.165.004).

**TwinsUK: St Thomas’ UK Adult Twin Registry**

The Twins UK study was funded by the Wellcome Trust, European Community's Seventh Framework Program (FP7/2007-2013)/grant agreement HEALTH-F2-2008-201865-GEFOS and (FP7/2007-2013), ENGAGE project grant agreement HEALTH-F4-2007-201413, and the FP-5 GenomEUtwin Project (QLG2-CT-2002-01254). The Twins UK study also receives support from the Department of Health via the National Institute for Health Research (NIHR) comprehensive Biomedical Research Centre award to Guy's and St. Thomas' NHS Foundation Trust in partnership with King's College London. TDS is an NIHR Senior Investigator. The Twins UK study also received support from a Biotechnology and Biological Sciences Research Council (BBSRC) project grant (G20234) and a U.S. National Institutes of Health (NIH)/National Eye Institute (NEI) grant (1RO1EY018246), and genotyping was supported by the NIH Center for Inherited Disease Research. The Twins UK study also received support from the National Institute for Health Research (NIHR) comprehensive Biomedical Research Centre award to Guy's and St. Thomas' National Health Service Foundation Trust partnering with King's College London.

**UK Biobank**

This research has also been conducted using the UK Biobank Resource under Application Numbers 11425, 12514 and 9797. Informed consent was obtained from UK Biobank subjects.

**UKHLS: Understanding Society – The UK Household Longitudinal Study**

The UK Household Longitudinal Study, led by the Institute for Social and Economic Research at the University of Essex is funded by the Economic and Social Research Council (Grant Number: ES/M008592/1). Data were collected by NatCen and the genome wide scan data were analysed by the Wellcome Trust Sanger Institute. Access the data at <https://www.understandingsociety.ac.uk/>

**WGHS: Women’s Genome Health Study**

The WGHS is supported by the National Heart, Lung, and Blood Institute (HL043851,

HL080467, HL09935) and the National Cancer Institute (CA047988 and UM1CA182913) with collaborative scientific support and funding for genotyping provided by Amgen.

**WLS: Wisconsin Longitudinal Study**

This research uses data from the Wisconsin Longitudinal Study (WLS) of the University of Wisconsin-Madison. Since 1991, the WLS has been supported principally by the National Institute on Aging (AG-9775, AG-21079, AG-033285, and AG-041868, R01 AG041868-01A1), with additional support from the Vilas Estate Trust, the National Science Foundation, the Spencer Foundation, and the Graduate School of the University of Wisconsin-Madison. Since 1992, data have been collected by the University of Wisconsin Survey Center. The opinions expressed herein are those of the authors. A public use file of data from the Wisconsin Longitudinal Study is available from the Wisconsin Longitudinal Study, University of Wisconsin-Madison, 1180 Observatory Drive, Madison, Wisconsin 53706 and at [http://www.ssc.wisc.edu/*WLS*research/data/](http://www.ssc.wisc.edu/WLSresearch/data/)

**QIMR: Queensland Institute of Medical Research**

Funding was provided by the Australian National Health and Medical Research Council (241944, 339462, 389927, 389875, 389891, 389892, 389938, 442915, 442981, 496739, 552485, 552498), the Australian Research Council (A7960034, A79906588, A79801419, DP0770096, DP0212016, DP0343921), the FP-5 GenomEUtwin Project (QLG2-CT-2002-01254), and the U.S. National Institutes of Health (NIH grants AA07535, AA10248, AA13320, AA13321, AA13326, AA14041, DA12854, MH66206). A portion of the genotyping on which the QIMR study was based (Illumina 370K scans) was carried out at the Center for Inherited Disease Research, Baltimore (CIDR), through an access award to the authors‘ late colleague Dr. Richard Todd (Psychiatry, Washington University School of Medicine, St Louis). S.E.M., is supported by an Australian National Health and Medical Research Council Fellowship APP1103623. The funders had no role in study design, data collection and analysis, decision to publish, or preparation of the manuscript. Researchers interested in using QIMR data can contact Nick Martin.
